# Correctness is its own reward: bootstrapping error signals in self-guided reinforcement learning

**DOI:** 10.1101/2025.07.18.665446

**Authors:** Ziyi Gong, Fabiola Duarte, Richard Mooney, John Pearson

## Abstract

Reinforcement learning (RL) offers a compelling account of how agents learn complex behaviors by trial and error, yet RL is predicated on the existence of a reward function provided by the agent’s environment. By contrast, many skills are learned without external guidance, posing a challenge to RL’s ability to account for self-directed learning. For instance, juvenile male zebra finches first memorize and then train themselves to reproduce the song of an adult male tutor through extensive practice. This process is believed to be guided by an internally computed assessment of performance quality, though the mechanism and development of this signal remain unknown. Here, we propose that, contrary to prevailing assumptions, tutor song memorization and performance assessment are subserved by the same neural circuit, one trained to predictively cancel tutor song. To test this hypothesis, we built models of a local forebrain circuit that uses contextual premotor signals to cancel tutor song auditory input via synaptic plasticity. After learning, excitatory projection neurons signaled mismatches between the tutor song and birds’ own performance, best matching experimental data when learning involved anti-Hebbian plasticity in recurrent interneurons. We also found that learning proceeds by both sharpening error sensitivity and minimizing responses to the tutor song. Finally, the error signals produced by this model can train a simple RL agent to replicate the spectrograms of adult bird songs. These results suggest that local learning via predictive cancellation suffices for bootstrapping error signals capable of guiding self-directed learning of natural behaviors.

**Significance:** Many species, including humans, learn new skills by trial and error. But in the absence of external rewards and punishments, individuals need a means of assessing the quality of their performance. Here, we model the process by which juvenile male zebra finches memorize the song of an adult tutor, showing how local changes in brain circuits responsible for hearing can give rise during this process to an internal error signal that later allows for self-guided learning.

---

The ability to acquire and maintain skilled motor behaviors is among the most impressive capabilities humans possess, from hitting a major league baseball to performing a Bach concerto. Yet learning these highly refined skills requires extensive practice, typically without the aid of external rewards or punishments. As part of this process, learners must therefore assess their own progress, presumably through a comparison of sensory feedback with some desired state. That is, for self-guided reinforcement learning to be possible, agents must somehow construct an internal reward function, a representation of high-quality performance capable of guiding subsequent behavioral adjustments.

In nature, the song copying behavior of male zebra finches provides a sophisticated example of precisely this type of learning. Early in development, juveniles memorize the courtship song of an adult male tutor during an initial “sensory” phase of learning, using it to guide independent practice in the subsequent “sensorimotor” phase, with the goal of replicating the tutor song (1). This copying process has typically been understood through the framework of reinforcement learning (RL) (2–5). In RL, animals alter their behaviors over time to gain rewards or avoid punishments, operationalized as maximizing the total reward delivered by the environment. In multiple species, such learning is known to be subserved by dopaminergic neurons in the ventral tegmental area (VTA), whose responses signal the mismatch between actual and expected reward (6–13). Likewise, dopamine has also been shown to be a critical driver for song learning in zebra finches (14–19), though in this case, it is an *internally-generated* reinforcement signal tracking performance quality relative to some standard (2, 4, 15, 18, 19). The fact that this standard — a particular tutor song — is not innate but learned thus raises the question of how juveniles bootstrap these performance errors.

Previous studies have implicated multiple interconnected auditory nuclei in signaling errors in birds’ own songs (20–22), including regions projecting to VTA (20, 21, 23, 24). Hypothetically, a neuron representing evaluative information should deviate from its baseline activity when sensory feedback during singing differs from desired behavior, while its activity should remain the same when performance meets expectations or the bird is silent. Indeed, certain neurons from Field L, caudolateral mesopallium (CLM) and ventral intermediate acropallium (Aiv) in adult birds respond to disruptive auditory feedback presented during singing but not to uninterrupted singing nor playback of normal or distorted song during non-singing periods (20, 21). Moreover, this error information is likely to converge in the intermediate arcopallium (Aiv) (21), since Aiv projects directly to VTA and pitch-dependent optogenetic activation of Aiv → VTA terminals results in negative reinforcement of syllable pitch in adult birds (23).

Another important finding is that several of these auditory regions directly or indirectly receive premotor inputs from area HVC (proper name) (21, 25–27). The presence of sensorimotor inputs to sensory regions suggests the possibility of corollary discharge mechanisms for the cancellation of self-generated sensory input, similar to those observed in mice (28–30), primates (31), and weakly electric fish (32–34). Likewise, there is evidence in adult birds that premotor input from HVC to auditory regions may govern the selective engagement of error-responding neurons during singing (4, 21, 23). Interestingly, HVC neurons have been found to exhibit tutor-song responsiveness in awake birds at early ages (35–40), a feature which declines with age (35) and disappears in adults (41, 42). Thus, both the sensory and premotor information necessary for computing performance errors are present directly upstream of VTA, though the mechanism by which vocal performance error is computed remains unknown.

Indeed, few studies have considered the problem of bootstrapping learning during the sensory phase. A previous theoretical proposal (43) posited that local learning rules could suffice for learning both forward and inverse vocal models without the need for an explicit template or reinforcement learning, but this required a specialized learning rule and an unknown gating mechanism that would selectively suppress recurrent auditory connections during tutoring. By contrast, we show below how a much simpler setup based on known physiology and simple Hebbian mechanisms can achieve the same “template-free” effect, producing error codes capable of guiding reinforcement learning during the sensorimotor phase.

Here, inspired by these corollary discharge ideas, we hypothesize that during the sensory phase of learning, local learning rules in auditory areas of juvenile finches facilitate predictive cancellation of a copy of tutor song using cues provided by premotor inputs. That is, we propose that tutor song memorization and performance error computation are not separate processes but subserved by a single circuit mechanism. As a result, projection neurons in these areas gradually form a sparse population error code sufficient for guiding reinforcement learning in the sensorimotor phase (Fig. 1A). To test this possibility, we implemented a set of models varying in circuit structure, locus of plasticity, and type of learning rule, comparing against existing data recorded in auditory nuclei. We found that a balanced excitatory-inhibitory network with anti-Hebbian plasticity in the recurrent connections involving interneurons was most consistent with experimental data. Moreover, in analyzing the learning dynamics of this network, we found that the “error landscape” — the geometry of population responses — is altered in two ways during the sensory learning period: First, the gain of error responses increases, sharpening the error landscape. Second, the minimum of this landscape gradually moves to align with the tutor song. Lastly, we found that, using the error codes acquired by fitting to real zebra finch song data, we could train a simple agent to accurately copy a tutor song spectrogram via reinforcement learning. Together, these findings show that purely local learning mechanisms for predictive cancellation are sufficient for estimating multidimensional performance errors, signals sufficient to enable self-guided learning of natural behaviors.

**Fig. 1.**
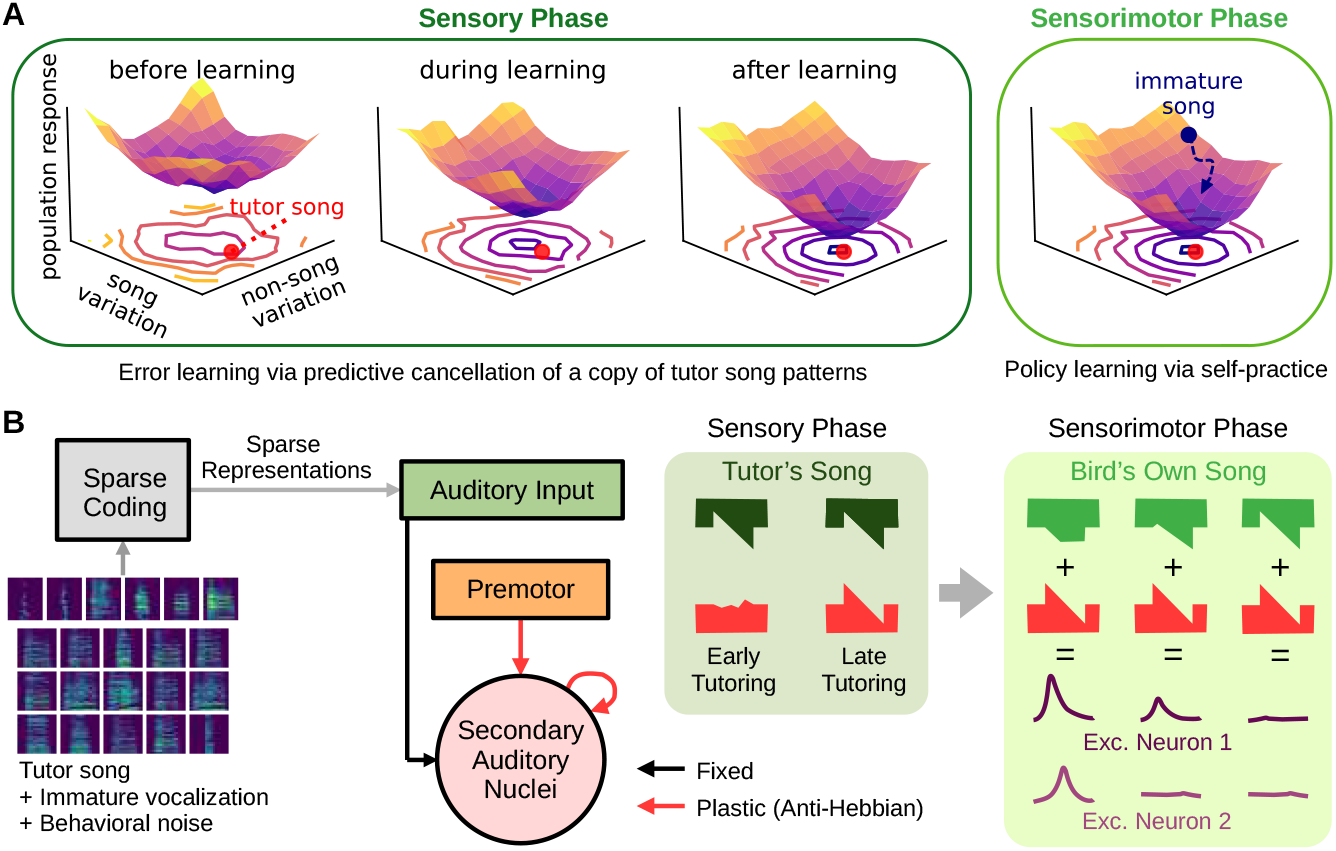
Learning to cancel tutor song results in an error landscape sufficient for subsequent motor learning. **(A)** Schematic of the general hypothesis. Neuronal responses to sensory input define a multidimensional, time-dependent “error landscape.” During the sensory learning phase, local learning rules increase the landscape’s curvature (sensitivity to error) and move its minimum toward the target tutor song (red dot). In the subsequent sensorimotor learning phase, this error serves as a negative reward, and reinforcement learning algorithms refine the bird’s own song by performing error minimization on this landscape. **(B)** General model structure. A sparse coding model maps tutor song to auditory input, which is sent on to meet song-locked premotor timing input in secondary auditory nuclei. Local plasticity rules in these regions gradually learn to perform predictive cancellation of this pattern, giving rise to sparse error codes.

## Results

To test for the emergence of error codes via local learning, we implemented a simplified model of the interconnected auditory areas of the forebrain as a single network receiving two inputs: upstream auditory input and premotor input (Fig. 1B). For the upstream auditory input, we assumed a sparse encoding of the song spectrogram (44, 45) and trained a classic sparse coding model (46) using a combination of adult male zebra finch songs (47), immature vocalizations (48), and noise caused by behaviors such as flapping. The outputs of this model were then used as the firing rates of the auditory input population (Fig. S1; see SI Methods).

For premotor input, we assumed a sparse, sequential activity pattern in which individual units burst at well-defined moments during the song, similar to observed activity in HVC (49, 50). While this pattern is well-documented in adult finches, it has also been observed that HVC contains tutor-song-selective neurons at early ages (35–38, 40). However, to test the sensitivity of our results to the precision of this premotor sequence, we also replicated our experiments using a “developing” premotor input in which the peak time and peak width of premotor input are initially strongly irregular and only gradually become regular over learning. We found that using such premotor input for training does not affect the ability of the network to learn error codes that match experiments (Fig. S2; see below).

We modeled the local auditory circuit as consisting of a single population of excitatory projection neurons receiving excitatory premotor input and both excitatory and inhibitory sensory input from the sparse coding network. In other models, we added to this excitatory population a reciprocally connected pool of local inhibitory interneurons. To test the effect of different types of local learning rules and synaptic loci, we considered four models: (1) A feedforward network of excitatory neurons, with anti-Hebbian plasticity in premotor →E connections (Fig. 2A) and three balanced excitation-inhibition (EI) networks of interconnected excitatory (E) and inhibitory (I) neurons (Fig. 2B) with differing loci of learning: (2) anti-Hebbian premotor →E connections; (3) anti-Hebbian E → E connections; or (4) *Hebbian* E →I and I → E connections^†^. That is, in Hebbian plasticity models, a synapse was strengthened (or weakened) if the pre- and postsynaptic neurons were co-active (or not co-active) (Fig. 2C), while for anti-Hebbian models, the contingency was reversed. For simplicity, we implemented this learning using bilinear plasticity rules (51).

**Fig. 2.**
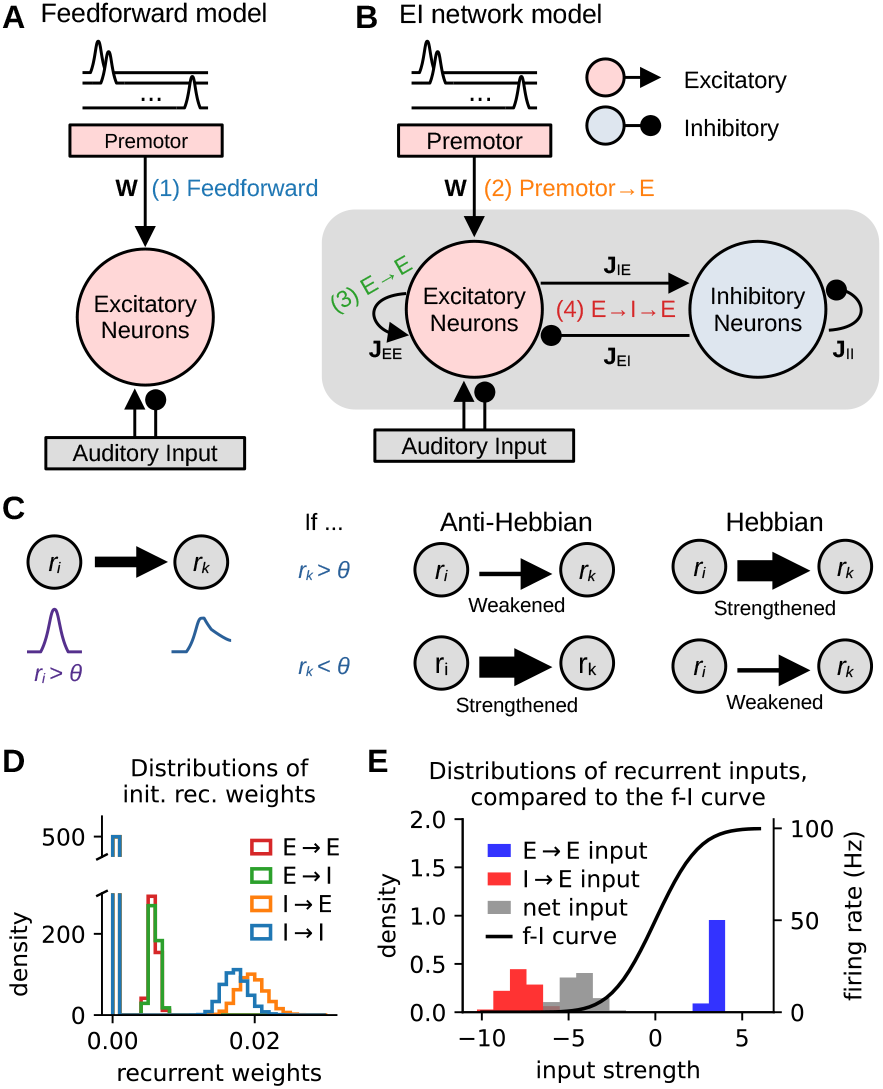
Four classes of models for local learning. **(A)** Feedforward model. Song-related premotor input directly drives excitatory neurons in secondary auditory areas, which receive both excitatory and inhibitory input from primary auditory areas. Plasticity takes place at the premotor→secondary auditory synapse (1). **(B)** Balanced excitation-inhibition (EI) networks with three potential sites of plasticity (2–4). As in **(A)**, excitatory neurons in secondary auditory areas receive input from both primary auditory areas and premotor areas, as well as a local population of inhibitory interneurons. Numbers indicate the site(s) of plasticity in each of the three models. **(C)** Schematic of rate-based Hebbian and anti-Hebbian plasticity rules. Hebbian rules strengthen synaptic connections when postsynaptic excitation exceeds a threshold *θ*. Anti-Hebbian rules do the reverse. **(D)** Distributions of initial recurrent weights for the four types of connections. The bins at zero represent the 50% of connections in each model type that are set to 0 for sparsity. In general, inhibitory weights need to be stronger than excitatory weights to achieve EI balance. A detailed description of the parameters and construction of weights is in Methods and Table 1. **(E)** Illustration of EI balance. Left vertical axis: density of the distributions of E→E (blue), I→E (red), and their net inputs (gray); right vertical axis: output firing rate of the f-I curve (black). EI balance is achieved when the distribution of net inputs lies near the activation threshold of the f-I curve.

To achieve excitatory-inhibitory balance, we initialized the recurrent connection weights of the EI networks to be strong and random, with means obeying the balanced condition (52) (Fig. 2D), such that the net recurrent input to a neuron lay within the linear regime of its activation function, even if recurrent excitation or inhibition would separately have produced runaway or silent activity, respectively (Fig. 2E). Together, these models produced sparse activity at biologically plausible firing rates while allowing us to consider multiple possibilities for both circuit connectivity and the locus of synaptic plasticity during learning.

**Table 1.**
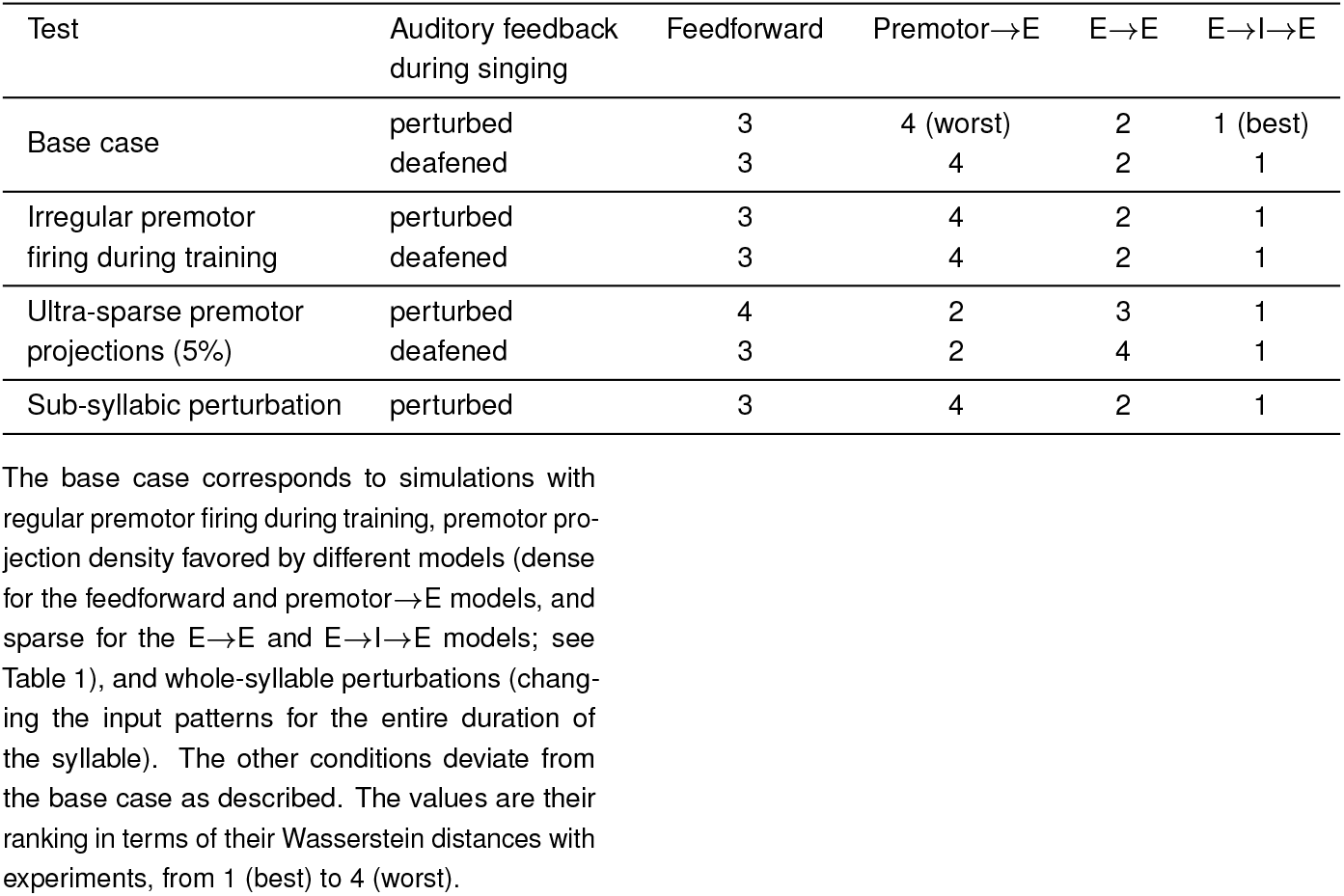
Summary of comparisons between models and experiments.

### Learning predictive cancellation enables neurons to signal error

We simulated the sensory phase of learning by training each of our models to cancel sparse auditory embeddings of the spectrograms of real adult male zebra finch songs (Fig. 1B). As expected, anti-Hebbian learning reduced the activity at both the population and individual neuron levels (Fig. 3A-B), and plastic weights projecting to the excitatory neurons of auditory regions were more negatively correlated with tutor song input after learning (Fig. S3), allowing local input to cancel the expected auditory pattern of the tutor songs.

**Fig. 3.**
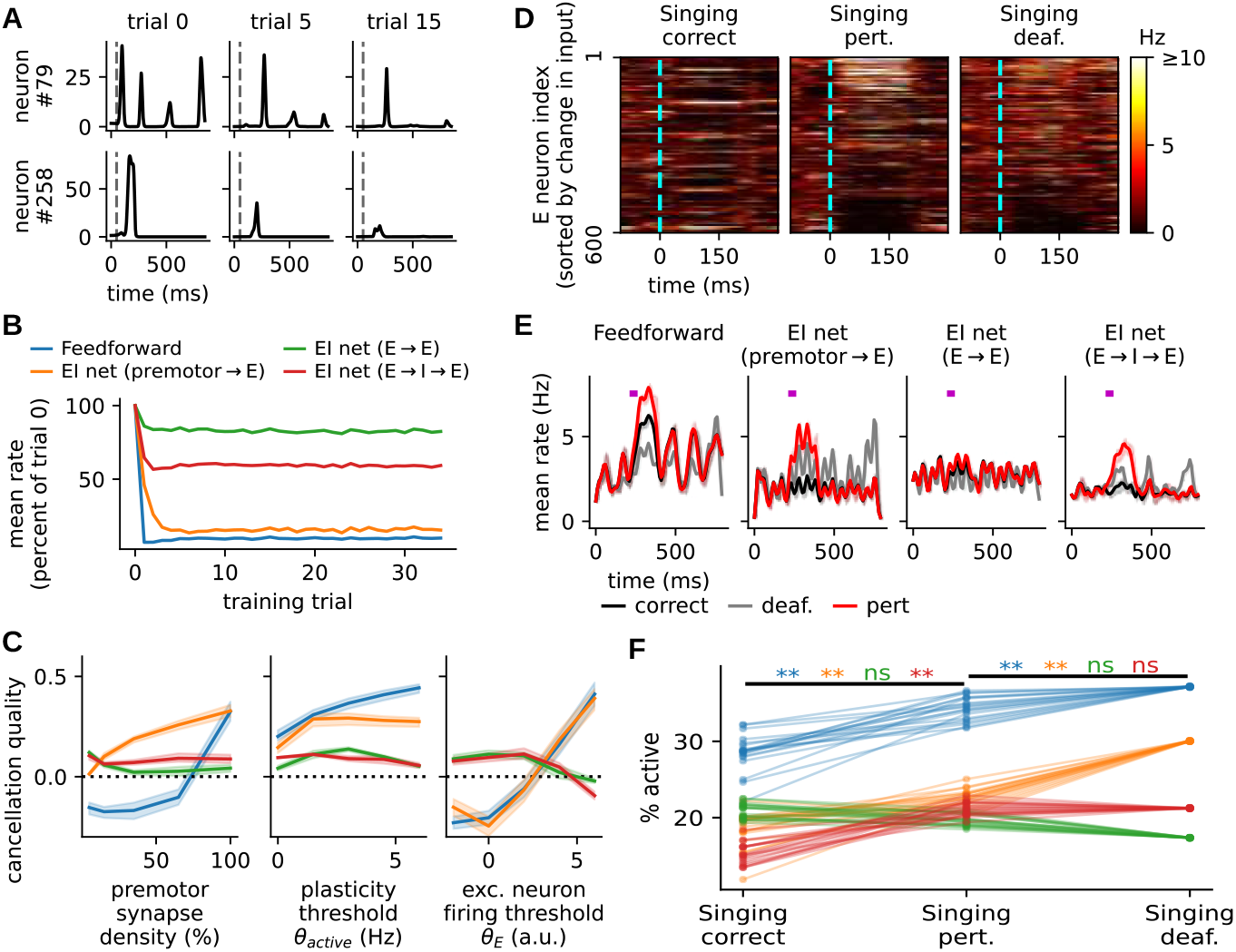
Neurons produce sparse error codes after learning to cancel tutor song auditory patterns. **(A)** Two example neurons in the E→I→E model decrease firing over learning. Gray dashed vertical lines mark song onset. Other models are qualitatively similar. **(B)** Population mean firing rates decrease over training, relative to the initial population means. Feedforward and non-recurrent models exhibit the largest percent decreases. **(C)** Quality of cancellation (see SI Methods) in the four models as functions of premotor projection density, plasticity threshold *θ*_active_, and excitatory neuron activation threshold *θ_E_*. Positive (or negative) quality of cancellation indicates lower (or higher) similarity of population activity with tutor song patterns after training. Widths of the shaded areas denote 1 s.d. across *n* = 5 initializations of each model. Models with recurrent plasticity perform best in the physiological regime of sparse inputs and low firing thresholds. **(D)** Learned singing responses under practice and perturbation. Raster plots show the responses of all excitatory neurons during singing under conditions of correct song (matching the tutor song), perturbation (added white noise), and deafening (no auditory input). Neurons are sorted by the difference between auditory input (bird’s own song) patterns and the sample-averaged tutor song pattern. **(E)** Population mean rates in the correct (black), deafened (grey), and perturbation (solid red) cases during singing. The purple bar in each subplot indicates the 50-ms white noise perturbation to the auditory feedback of the bird’s own song. Only the E→E model fails to exhibit a population response to error. **(F)** Percentages of active excitatory neurons (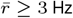, see SI Methods) in different models, averaged across all samples, from the earliest perturbation onset to 100 ms after the latest perturbation offset. Except for the E→E model, a significantly denser set of neurons are active during perturbed singing than during correct singing (double asterisks indicate *p* < 10^−3^, one-sided permutation test). Note that the learning rate is set to zero in **(D-F)**.

Comparing across models, we found that while firing rate reductions were more pronounced in the feedforward and premotor →E models (Fig. 3B), the quality of tutor song cancellation depended on multiple parameters. In particular, we probed how the cancellation depended on three important and experimentally measurable quantities: the density of premotor synapses, the equilibrium threshold of plasticity, and the excitatory neuron firing threshold (Fig. 3C). When premotor projection density is sparse, as observed in experiments (21, 27), cancellation is more effective in the E→E and E→I→E models than the models with premotor →E plasticity, because the latter models must directly store negative images of tutor song patterns in the limited premotor →E weights. In this sparse premotor projection scenario, the error codes in the feedforward and premotor →E models are degraded (Fig. S4). Furthermore, the quality of cancellation remains relatively stable over different plasticity thresholds across all models, while for the excitatory neuron firing threshold, the feedforward and premotor →E models fail to cancel tutor song patterns unless the excitatory neurons are hard to activate, while the E→E and E→I→E models favor intermediate to low thresholds (Fig. 3C). Thus, models with recurrent plasticity are more robust to different parameter values in our models.

Having verified that models could reliably cancel tutor song, we then proceeded to assess their error-coding capacity by testing their responses to unanticipated auditory inputs. Following experiments in which the responses of auditory neurons were probed by playing either undistorted or noise-corrupted recordings of the bird’s own song (20, 21), we set the learning rates of our fully-trained models to zero and observed their responses under these conditions. To simulate the correct singing case, in which the bird’s own song matches the tutor song, we randomly selected samples from tutor song data as the auditory inputs. In response to these stimuli, most excitatory neurons in a local forebrain circuit exhibited low transient activity (Fig. 3D). In contrast, when the auditory feedback of the bird’s own song was perturbed by directly adding 50 ms of white noise to the stimulus, these neurons displayed heterogeneous error responses, with larger firing rates on trials with larger mismatch between auditory feedback and the sample-averaged tutor song. In addition, when the auditory feedback was removed, simulating the case of singing in deafened birds, neurons weakly signaled the *inverse* pattern of tutor song—indicative of predictive cancellation. At the population level, the mean rates (Fig. 3E) and percentages of active neurons within the perturbation time window (Fig. 3F) of the excitatory population were increased by perturbation and deafening in all but the E→E model. Though the E→E model did not exhibit a mean population error response, it nonetheless showed heterogeneous error responses like the others (Fig. S5A-D).

We also observed that increases in population mean firing rates and percentages of active neurons did not persist to the end of the song (Fig. 3E and Fig. S5E), agreeing with experimental findings (21). Using input activity perturbations (analogous to stimulating the axon terminals from upstream auditory regions to these secondary auditory nuclei) much shorter than the duration of a syllable, we further verified that the circuits can signal error on a sub-syllabic basis (Fig. S6A-C). Together, these results suggest that, simply by using local learning to cancel tutor song, our models simultaneously developed sparse population error codes with high temporal precision.

### Experimental data favor error codes learned via inhibitory plasticity

As stated above, our models considered four potential kinds of local plasticity in auditory circuits. To determine which of these candidate model is most likely, we compared the error codes from our models with neural activity obtained from one-photon calcium imaging performed in the lateral caudal mesopallium (CML) of adult male zebra finches (Fig. 4A) singing undirected song with white noise perturbation before and after deafening (Fig. 4B, n=8 birds, 397 neurons for white noise perturbation, n=6 birds, 161 neurons for pre and pos-deafening). CML is a primary auditory region and also receives input directly from the premotor area HVC and NIf. During recording, we used a closed-loop system (53) to randomly target 50% of syllable renditions with white noise to induce an artificial vocal error. We found that CM neurons displayed sequential activation or suppression during singing, with this activity disrupted by white noise perturbation (Fig. S7A). In a sparse set of neurons, mean responses to these perturbations increased relative to correct singing, while a larger subset of neurons increased their responses during song post-deafening (54) (Fig. 4B). Further, the mean responses of a few neurons strongly decreased during song post-deafening, but this was not observed in the white noise perturbation cases.

**Fig. 4.**
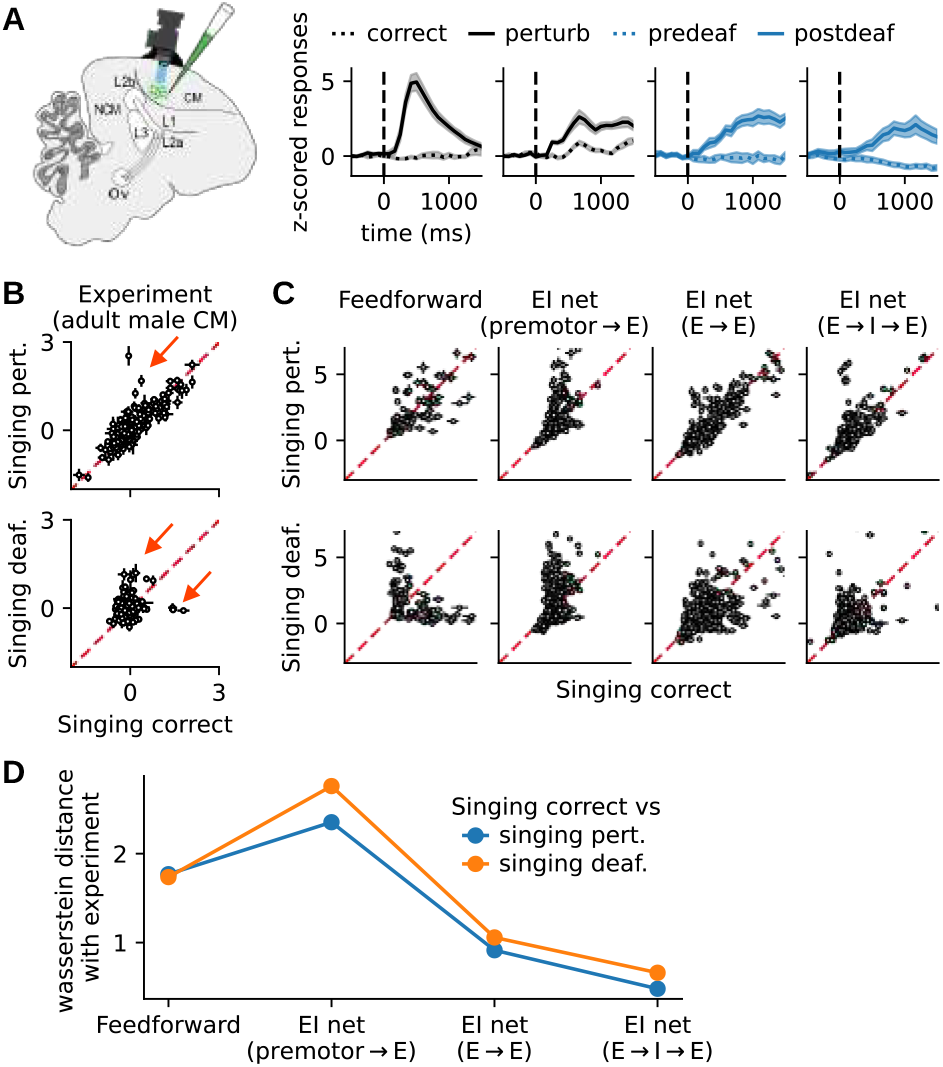
Comparison of experimental data to models with different sites of local plasticity. **(A)** Left: schematic of the pan-neuronal calcium imaging experiments. Right: z-scored responses of four example units in the case of normal singing (dashed lines) and singing with auditory perturbation or deafening (solid lines). **(B)** Distributions of trial-averaged z-scored calcium fluorescence in correct versus perturbed singing in area CM of adult male zebra finches. Top row: scatterplot of white noise perturbation responses versus responses on correct singing trials (*n* = 397 ROIs). A sparse set of neurons are more activated by white noise perturbation (red arrow). Each dot represents the trial average of the normalized activity of one neuron over the song period (see SI Methods). The horizontal and vertical error bars at each dot represent the standard errors for the correct and perturbed or deafened trials. Bottom row: same as top row, but comparing responses post-deafening with correct singing responses pre-deafening (*n* = 161 ROIs). **(C)** Same as **(B)** for the four classes of plasticity models (with model firing rates instead of fluorescence). Only 300 randomly selected neurons were plotted for visualization purpose. **(D)** Wasserstein distances between the distributions from experimental data **(B)** and from models **(C)** indicate that the E→I→E model is the closest to experimental data.

Our models replicate these effects. We found both intact and disrupted sequential activation during normal and perturbed singing, respectively, though only the E→E and E→I→E models could additionally produce the observed sequential suppression (Fig. S7B). Moreover, in comparing perturbed to unperturbed responses, we found a sparse set of neurons more activated by white noise perturbation and deafening than during normal singing, and such activation is stronger and denser in the deafening case than white noise perturbation (Fig. 4C).

Yet, while all models contained such a sparse population of perturbation-responsive neurons, not all models provided an equally good quantitative match to data. To quantify the prevalence and magnitude of this perturbation response across models, we measured the Wasserstein distances between the correct-perturbed or correct-deafened joint distributions from experiments and model simulations (Fig. 4D; see SI Methods). The lower this distance, the closer the models are to the experiments. We found that models with recurrent plasticity matched experimental data better than models with plasticity in the feedforward synapses from the premotor area, and the E→I→E model matches experiments the best among all four models. This observation still holds in more realistic and thus more challenging settings, such as training with initially irregular but gradually organizing premotor activity (Fig. S2), training and testing with sparse premotor synapses (Fig. S4), and testing models with perturbations that only cause short (50 ms) changes of input firing patterns (Fig. S6). A brief summary of the results of these simulations is provided in Table 1, which suggests that the E→I→E model with local inhibitory plasticity best matches experimental data.

### Population error codes exhibit two-stage learning

In our models, local learning rules give rise to predictive cancellation of tutor song inputs, producing as a byproduct sparse population error codes. Viewed as a function of auditory input, these codes describe an “error landscape” whose geometry might guide subsequent reinforcement learning. In principle, a good error landscape should produce the lowest error for all variants of the correct song (the manifold of correct song), and be sensitive to songs deviating from this manifold. To draw intuition about how the error landscape evolves during training, we periodically froze the networks during training and recorded the singing responses when auditory feedback was perturbed (Fig. 5A). Specifically, we considered two types of perturbations: Perturbations in the “off-manifold” direction interpolate between feedback generated by the correct song pattern and a completely random pattern, while perturbations in the “on-manifold” direction interpolate between the correct singing (1) and deafened/no feedback (0) cases. We found that the slope of the population response rapidly grows in the off-manifold direction over training, suggesting a strong increase in the sensitivity to different auditory patterns during singing (Fig. 5B). Along the on-manifold direction, learning shifts the auditory feedback eliciting minimum singing response away from silence and toward to the tutor song pattern (Fig. 5C). Thus, learning alters both the sensitivity and selectivity of auditory feedback during training.

**Fig. 5.**
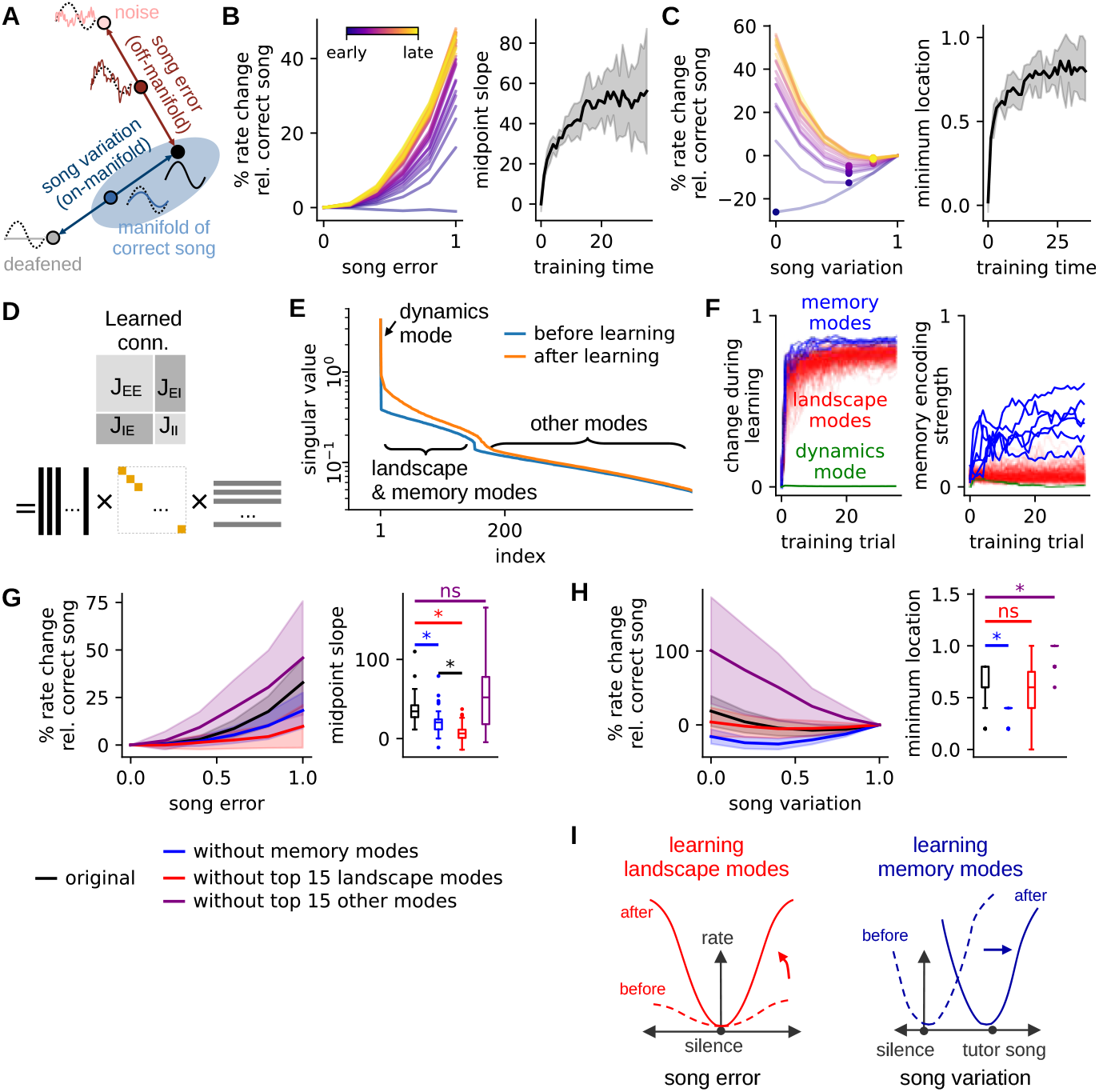
The minimum and shape of the error landscape are encoded by different connectivity modes in the E→I→E model. **(A)** Schematic of acoustic variation used to probe learning. Auditory feedback inputs were varied both in the song error direction (dark red arrow), which interpolates between the correct song and a random pattern, and song variation direction (dark blue arrow), which scales the correct song pattern between deafened and correct singing. The song error direction represents auditory input off the manifold of correct song (light blue region), while the song variation direction partially overlaps with the manifold. **(B)** Left: Mean relative singing responses to the auditory feedback in (B) the song error direction during training. 0: correct song; 1: random auditory feedback pattern. Relative responses are defined as the percent change in population firing rate compared to the correct singing responses. Learning progress increases from purple to yellow. Right: The midpoint slope is the slope of the line tangent to the curves at 0.5 on the left plot. Higher values indicate greater sensitivity to auditory feedback. Widths of the shaded areas denote 1 s.d. across *n* = 10 simulations. **(C)** Left: Mean relative singing responses to the auditory feedback in the song variation direction during training. 0: no auditory feedback (deafened); 1: correct song. The dot on each curve in the left panel marks its minimum. Right: Minimum location of the responses in the left panel. Higher values indicate minimal error responses to away from silence and closer to the tutor song. **(D)** Schematic of the singular value decomposition of the concatenated connectivity matrix. **(E)** The sorted singular value spectra before (blue) and after (orange) learning. The spectrum exhibits a clear kink that we use to differentiate between dynamics, landscape, and memory modes and “other modes.” **(F)** Left: Dissimilarity between the non-memory modes and their initial values over training time. The green curve is the dynamic mode, which barely changes, and the red curves are landscape modes, which change rapidly and quickly stabilize. Right: Strength of memory encoding for all modes during training. The blue curves are those with final memory encoding strengths larger than the maximum pre-learning memory encoding strength (i.e., noise correlation). **(G, H)** Altering landscape and memory modes alters population error responses. Comparisons among the original trained models (black); models with the top 15 landscape modes removed (red), with memory modes removed (blue); and with the top 15 non-memory, non-landscape modes removed (purple). Widths of the shaded areas denote 1 s.d. across *n* = 35 simulations of each model. Asterisks in the the right panels indicate significant differences between distributions (*p* < 0.01; two-sided Wilcoxon signed-rank test; colored asterisks: altered model versus original model; black asterisks: models with perturbed memory versus landscape modes). Perturbations of the landscape modes primarily affect the error landscape slope, while perturbations of the memory modes alter its minimum. **(I)** Schematic of changes to the error landscape as a result of learning.

To further understand how the error code emerges in the dynamics, we analyzed the spectrum of the recurrent weight matrix **J** of our most plausible candidate, the E→I→E model, over the course of training. Since this matrix governs recurrent population dynamics in response to auditory input, understanding its changes during learning sheds light on the process by which the error landscape takes shape. Specifically, we analyzed **J** using its singular value decomposition (SVD), decomposing it into a sum of rank-1 input-output pattern pairs (Fig. 5D-F). Each of these pairs describes a “singular mode,” an independent pattern of activity shaping the population responses.

We found that, while **J** remains high-rank throughout learning (Fig. 5E), its singular modes are readily separable into groups based on their learning time courses and effects on the error landscape. In all cases, the top singular mode of **J** was unchanged by learning (Fig. 5E-F) and encoded the network’s basic connectivity structures and responses to transients; perturbing it resulted in runaway dynamics (Fig. S8). Since this mode ensures the dynamic balance of neural activity, we refer to it as the “dynamic mode.” After the dynamic mode, we found range of singular modes that changed significantly over the course of learning (Fig. 5E-F). Among these, a small number (*N* ≤ 15) eventually developed correlations with tutor song patterns, while a second group that did not exhibit this pattern nonetheless changed rapidly and quickly stabilized after learning began. For reasons that will become clear below, we will refer to these as groups as “memory modes” and “landscape modes,” respectively. Finally, beyond these groups remained a long tail of modes (“other modes”) with much smaller singular values that we did not analyze.

How do these distinct classes of modes affect the learned error landscape? To understand this phenomenon, we performed perturbation experiments in which we altered one or both of sets of modes in the network weights post-learning. Specifically, we either (1) “de-memorized” the models by removing the correlation between memory modes and tutor song patterns^*^, or (2) shuffled the entries of a similar number of landscape modes or (3) shuffled other nonessential modes (see SI Methods). All three interventions significantly changed the mean firing rate of the network compared to the original models in the unperturbed, deafened, and perturbed singing cases (*p* < 10^−4^, two-sided Wilcoxon rank-sum test; Fig. S9A). To interpret the effects of intervention on the error landscape, we then tested the altered networks under the on- and off-manifold alteration of the auditory feedback introduced earlier (Fig. 5A-C).

We found that disrupting the landscape modes strongly flattened the error landscape (Fig. 5G) and elevated error responses (Fig. S9), indicating that the landscape modes primarily alter the curvature of the error landscape. On the other hand, decorrelating the memory modes from tutor song moved the minimum location of the error landscape away from the tutor song and closer to silence (Fig. 5H), suggesting that the role of these modes is to shift the minimum of the error landscape. Paradoxically, disrupting the “other modes” could even slightly increased the slope of the error landscape and moved the minimum location closer to the original tutor song pattern (Fig. 5G-H). This suggests either that some memorization of the training data has occurred in these small modes (55) or that local learning has not fully converged at the fixed step size chosen in our experiments. These observations are qualitatively similar across alternative shuffling methods (Fig. S10).

Together, these results suggest that changes in the recurrent connectivity matrix of the E→I→E model over the course of learning proceed in two ways: First, rapid learning of the landscape modes increases the gain of error responses and thereby sharpening the error landscape. Second, a relatively slower lock-in of memory modes aligns the minimum of the error landscape (smallest error response) with the tutor song pattern, providing a target for subsequent sensorimotor learning (Fig. 5I).

### Model error codes suffice for reinforcement learning of song

In the previous sections, we have shown that our models, trained with local learning rules, produce sparse population error codes and that the learning process consists of an initial phase of error gain adjustment followed by a shift in minimum error response toward the tutor song. However, it remains to be seen whether these error codes contain sufficient information about deviations between vocal performance and the tutor song to guide learning. To test this, we developed a simple learning paradigm based on the actor-critic RL framework, widely used to model vocal learning in songbirds (4, 23, 24, 56–58).

We assume that the evaluation of song generation and learning happen at the syllable level. For every syllable index (“state”) *s*(*t*), the actor generates a set of weights **a**(*s*) used to linearly combine a set of spectrogram basis elements stored in the rows of a matrix **B**. That is, the predicted spectrogram is **a**(*s*)^⊤^**B** (Fig. 6A). As above, the generated spectrograms undergo a sparse linear embedding, forming the auditory population input to our local circuit model. Rather than maximizing external rewards from the environment, our agent is trained to minimize the population averaged error signal at each time step, equivalent to maximizing an internal reward signal equal to the negative of this quantity: 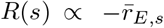, where 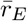 is the mean response of the excitatory population. The agent’s actions are parameterized as a state-dependent Gaussian distribution over basis elements: **a**(*s*) ∼ 𝒩(***µ***(*s*), **I**), and at each step of learning, the temporal difference (TD) error signal *δ* is used to update the agent’s estimate of the value of each state *V* (*s*). Specifically, after each syllable, these functions are updated by a standard temporal difference learning algorithm:

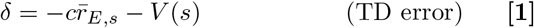

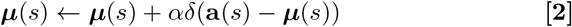

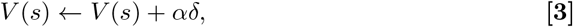

where *α* is the learning rate and *c* = 10 is a constant scaling the mean rate.

**Fig. 6.**
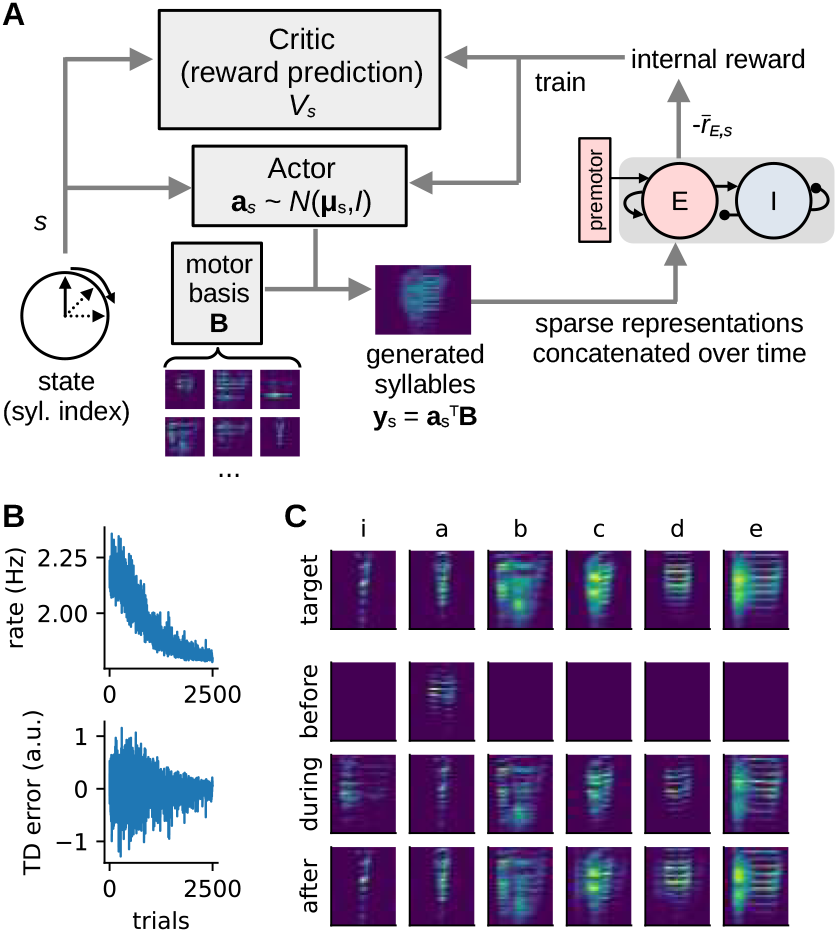
Error codes produced by local learning rules can be used to train a motor policy. **(A)** Schematic of the actor-critic reinforcement learning model. A syllable-based state variable *s*(*t*) indexes both the motor policy (actor **a**(*s*)) and the prediction of the expected reward (critic *V* (*s*)). Syllables **y**(*s*) are generated by linearly combining spectral basis elements **B** using the outputs of the policy **a**(*s*). The negative of mean error signal 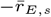 serves as the reward signal to be maximized. **(B)** Excitatory population firing rate (error signal) and temporal difference (TD) error plotted as a function of trial. Results shown are for the E→I→E model, but results for the feedforward and premotor→E models are similar (see Fig. S11). **(C)** Tutor syllable templates (top row), and model-produced song before, during, and after learning (last three rows). The model is able to reproduce the tutor song using only the learned population error code as feedback.

Using this learning model, we tested the ability of all four of our models to copy a tutor song using their own error signals as a negative intrinsic reward. Learning in this highly simplified task converged quickly, within 2500 trials, as indicated by the exponential decrease in vocal error and the convergence of the TD error to zero (Fig. 6B; Fig. S11A,C). For the E→E model, learning was slower, and only some generated syllables matched the target (Fig. S11D), likely because the population mean rates are largely insensitive to error (Fig. 3E). For the E→I→E (Fig. 6C), as well as the premotor →E (Fig. S11A) and feedforward models, all the generated syllables gradually matched the target syllables. Furthermore, when landscape and memory connectivity modes were perturbed as in our analysis of the previous section, learning in these perturbed models was impaired, with perturbed memory modes having the strongest effect (Fig. S12). Together, these results demonstrate that the models’ learned error codes are sufficient to guide a motor program to produce songs that resemble a tutor target.

## Discussion

For animals to learn complex behaviors without external reinforcement, they need an internal mechanism to evaluate their performance. Here, we have used computational models to explore the idea that this internal error is learned via local mechanisms orchestrating predictive cancellation of an expected sensory target. For any given sensory input, mismatches in this cancellation give rise to population error codes that can be used to move performance toward the target. Using songbird vocal learning as paradigmatic example, we implemented plausible circuit models with different connectivity motifs and types of plasticity that learn to cancel the auditory patterns of tutor songs during an early period of sensory learning. In the subsequent practice-based sensorimotor learning phase, auditory input to the system differs from the pattern of the tutor song, and an error code with similarities to observed responses in songbird auditory regions emerges. Among these models, the balanced excitation-inhibition network with Hebbian E→I and I→E plasticity rules (E→I→E model) best matched these experimental data. By characterizing changes to the weight matrix in the E→I→E model during learning, we also found that learning the cancellation proceeded in two ways: sharpening of error responses and reshaping of the geometry of population errors that shifted minimum firing away from silence and toward the target pattern. Finally, we confirmed that the error codes generated by the E→I→E model are capable of guiding a motor policy to produce songs that match the target using a simple reinforcement learning model.

To our knowledge, this work offers one of the earliest concrete computational models of *vocal error evaluation* in a local forebrain circuit in songbird auditory areas. Given the rich connectivity among these auditory areas (22, 59) and the fact that several sites are likely to be important to sensory learning (21–23, 27), we have treated them as a single balanced EI network model, assuming that the excitatory population both receives premotor information and sends error codes (21, 23, 25, 26). Based on findings that tutor-song-selective neurons are present in area HVC during early sensory learning (35–38, 40), we have also assumed that premotor input provides the temporal basis for predictive cancellation of tutor songs. While this idea of error codes arising from predictive cancellation is not new (e.g., 32, 33, in electric fish) and has been discussed in the context of novelty detection in the avian brain (60), ours is the first study to systematically examine both the implications of different types of plasticity in such a system and to demonstrate how the resulting error codes can subserve reinforcement learning.

Our model thus shares several features with a previous modeling study of song learning (43), which also proposed a form of local learning without an explicit representation of the tutor song. In that work, local plasticity in sensory areas during tutoring results in a predictive model of auditory activity evoked by tutor song. Our model differs from this one in three key ways: First, the focus of (43) was on inverse models as alternatives to reinforcement learning, while our focus is on learning error codes capable of guiding RL. Second, the model of (43) assumed a gating mechanism that selectively disabled recurrent sensory connections during the learning phase, a phenomenon that currently lacks empirical support. Our model, by contrast, focuses on predictive cancellation of sensory and premotor signals based on established physiology and requires no such gating mechanism. Third, the model of (43) posits plasticity based on synaptic competition and local eligibility traces for auditory activity, whereas ours requires only simple (anti-)Hebbian rules. Along related lines, another recent work (61) proposed a model capable of signaling error after predictive associative learning, which minimizes the gap between the actual firing and the prediction from recurrent excitation. When neurons receive external patterns, this learning rule effectively builds association between these patterns in the recurrent weights. On the contrary, our learning rules are anti-associative and produce signals representing the subtraction between sensory feedback and a learned predictive pattern. We demonstrated that this straightforward signal can be directly read out for RL.

Evidence from recent research offers some support for these results. Our model comparisons suggest that the E→I→E model is the most plausible (Fig. 4, Fig. S2), and studies have revealed age-contingent Hebbian plasticity from inhibitory interneurons to pyramidal neurons in mouse auditory cortex (62–64) that can affect both EI balance and sound frequency representation. Prediction error patterns similar to error codes in our models were also found in the single neuron responses in several auditory areas in anesthetized European starlings listening to conspecific songs (65), consistent with a predictive coding model. Other support comes from the zebra finch brain, where the premotor area HVC projects sparsely to secondary auditory areas such as CM and Aiv via avalanche (Av) (22, 59), though these projections may not be dense enough (21, 26, 27) to support the necessary learning in premotor →E models. However, the E→I→E model is robust to sparse premotor projections. In addition, spontaneous firing in Aiv (21) and CM (66) may indicate low neuronal firing thresholds, to which our E→I→E is also robust.

To confirm that the learned error codes are sufficient to guide song generation, we implemented an actor-critic RL system whose components align with prominent theories about learning in the avian song system (3, 4). For simplicity, we chose a scalar critic function and used the mean excitatory rates from our vocal error models to construct the temporal difference (TD) error, and under this assumption, the premotor →E and E→I→E models are capable of learning to produce all syllables, while the E→E model results in matches between some syllables. Yet, our models potentially support multidimensional error signaling, as suggested by the correlation between the error codes and the true difference between the target and actual input patterns, (Fig. 3D), as well as the fact that anti-Hebbian learning encodes negative templates in the connections (Fig. S3).

Consistent with the vector-valued error code observed in our models, stimulation of ventral pallidum (VP) and Aiv, leading candidates for the critic and vocal error estimator, respectively, drives heterogeneous responses in VTA (24). These heterogeneous responses have recently been linked to vector-valued dopamine signals (13, 67–69), for which several theories have been proposed. For example, heterogeneous dopamine responses have been suggested to encode prediction errors for different expectiles of the reward distribution (69), TD errors corresponding to subspaces of cortical states (67), or gradients from the mismatch error between basal ganglia outputs and targets (68). An equally important question is how these vector dopamine signals are constituted physiologically. Here, we have proposed biologically plausible models that support such vector error codes, and a critical future step will be to connect these models to the existing theories of vector dopamine feedback.

Lastly, while we have attempted to align our models with known zebra finch physiology, we have made several key simplifications for purposes of theoretical tractability and computational efficiency: First, auditory inputs to our model are temporally interpolated sequences of sparse embeddings of song syllable spectrograms (Fig. S1) produced by a classic linear sparse coding model (46, 70). This is, at best, a caricature of early auditory processing and discards sub-syllabic variation. Even so, anti-Hebbian learning of the type we propose is theoretically agnostic to input distribution as long as these inputs maintain reasonable variability. Thus, our models could, in principle, handle sub-syllabic structure by a proper choice of time constants and input scaling. Second, our models learn through bilinear Hebbian or anti-Hebbian rules with fixed thresholds (Fig. 2C and Eq. 6). In reality, the plasticity can be bounded, adaptive, and multi-factored. In this work, we have elided these complexities, focusing simply on what is learnable in principle by simple local rules. Third, we have focused on firing rates and rate-based neurons. Future studies will need to extend these results to the added sparsity and nonlinearities of spiking neural networks. Fourth, we do not presume to account for all auditory processing. While responses of our proposed cancellation circuit are minimal when the bird’s own song closely matches the tutor memory, we do not propose that the bird fails to perceive its own song in these cases. That is, the songbird’s predicament differs from other situations in which the goal of corollary discharge is to *perceptually* cancel predictable nuisance inputs (28, 29, 32).

In summary, our models suggest that local learning for predictive cancellation is sufficient for the formation of error signals that can guide self-driven learning of complex motor behaviors. Further, by comparing models with different types of plasticity and aligning these to experimental data, we predict that a circuit involving plasticity between excitatory and local inhibitory neurons is most likely to support learning of internal behavioral evaluation signals. Together, these results constitute a novel computational theory for the circuit mechanisms underlying animals’ ability to evaluate and refine their behaviors without external supervision or reinforcement.

## Methods

### Vocal error circuit models

The vocal error circuit models can be described by a set of differential equations for the total input 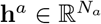 (where *a* = *E, I*)

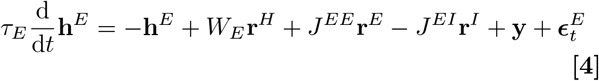

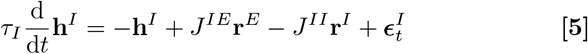

where **r**^*E*^ = *ϕ*_*E*_ (**h**^*E*^), **r**^*I*^ = *ϕ*_*E*_ (**h**^*I*^) and 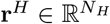 denote the excitatory, inhibitory, and premotor neuron firing rates, respectively, the *τ* ‘s are time constants, and

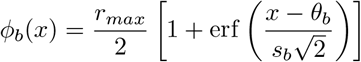

are sigmoidal activation functions. Values of the parameters for the main results are summarized in Table 1. Reasonable choices of maximum firing rates 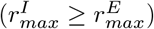 did not qualitatively affect the dynamics. *W*_*E*_ are the synaptic weights from premotor input to the excitatory neurons, *J*^*ab*^ ≥ 0 are local synaptic connections from population *b* to population *a*, **y** is the auditory input, and 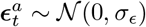 is the i.i.d. white noise at each time step.

### Hebbian and anti-Hebbian learning

In the premotor →E and E→E models, anti-Hebbian learning happens in either premotor →E or E→E connections. In the E→I→E model, Hebbian learning happens in E→I and I→E connections. Both learning rules can be described by the same differential equation, where the learning strength *η* is strictly positive for Hebbian learning, and strictly negative for anti-Hebbian learning.

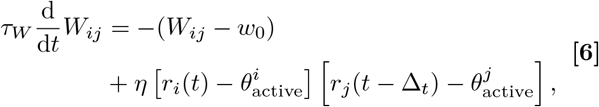

where *W*_*ij*_ is the synaptic weight from neuron *j* to neuron *i, τ*_*W*_ ≫ 0 is the time constant of the weight evolution, *w*_0_ is the weight baseline, Δ_*t*_ is the time asymmetry of the plasticity, and *θ*_active_ ∼*O*(1) are thresholds for determining whether the pre- or postsynaptic neurons are active. The first term is used to prevent the weights from growing to infinity. The second term learns the correlation or anti-correlation between the pre- and postsynaptic neurons. Briefly, in (anti-)Hebbian learning, the synaptic weight is increased (decreased) when both the pre- and postsynaptic neurons are active, and decreased (increased) when they are not co-active. To obey Dale’s law, the weights are clipped at each time step:

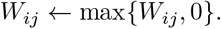

Values of the parameters are summarized in Table 1.

### Premotor Input

We assume that premotor neurons in our model respond to tutor song, as has been found for HVC (35–38, 40). Specifically, the firing rate of a premotor neuron 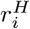 for *i* = 1, 2, …, *N*_*H*_ in multiple renditions *j* = 1, 2, …, *N*_rend_ is modeled as

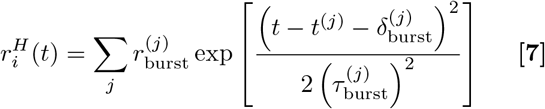

where 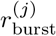 is the peak burst firing rate, 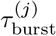 is the peak width, *t*^(*j*)^ is the expected burst time, and 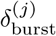 is the jittering for the burst peak. *t*^(*j*)^ is defined as 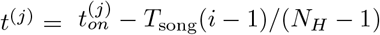, where 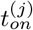 is the song onset, *T*_song_ is the song length. Notice that 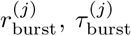, and 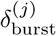 can vary over renditions.

Except for Fig. S2, we further assume that premotor neurons have already developed sparse bursting behaviors with stable burst times and relatively stable burst profiles during a rendition of song. We set 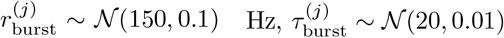, and 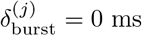.

We also tested the models using developing, progressively regular, premotor bursting profiles in Fig. S2. In these cases, 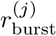 follows a log-normal distribution with mean 150 Hz and standard deviation 70*ψ*(*j*) Hz, 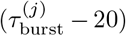 follows an exponential distribution with scale parameter 60*ψ*(*j*) ms, and 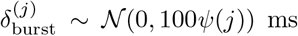, where

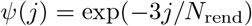

is a scaling factor decreasing smoothly over training.

### Auditory Input

Except for Fig.1A, we tested the models using input patterns derived from birdsong data. To simulate input to the secondary auditory regions, we postulate that the upstream auditory processing implements a form of sparse coding (70–72). We trained a sparse coding model (see Supplementary Methods) on flattened spectrograms of tutor song, similar to what a juvenile bird might hear during the sensory phase. More specifically, the training data involved the syllables of adult zebra finches, vocalizations of juvenile zebra finches (35-40 dph), and behavioral noise recorded during 35-40 dph, extracted and processed from (48) using the AVA library (73). After training, the sparse coding model trained using test sequences of syllable spectrograms, generating sequences of discrete syllable representations. To produce time series whose durations matched the original song recordings, each representation was extended over the time window of the corresponding syllable, with the gaps between filled by zeros. Lastly, the time series were convolved with an exponential kernel to smooth the transitions between representations.

### Perturbation and Deafening

To simulate external perturbation of auditory feedback (except for Fig.5F), we first added 50 ms of white noise to the audio clips of a particular syllable, generating the spectrograms using the same parameters as for unperturbed song. The spectrograms of both perturbed and unperturbed syllables are then fed into the sparse coding model in the same syllable order, and the continuous time series were generated following the same procedures as for the unperturbed input (Fig.S1). To simulate the deafening case, we set the auditory input **y** = 0.

To quantify the changes of error landscape in Fig. 5F, we directly perturbed the sparse embedding of the tutor song input because the alternative perturbation approach described above leads to great variance in the effective difference between perturbed and correct tutor song patterns for each choice of perturbation strength (Fig. S1G). First, a noise pattern, 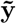 is generated by randomly shuffling the elements of the sparse embedding of the syllable to be perturbed. Given a perturbation strength variable *γ*, the new perturbed pattern, **ŷ**(*γ*) was calculated as

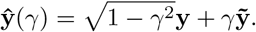

For Fig. 5G, we simply scaled the sparse embedding by a scaling parameter *a*: **ŷ**(*a*) = *a***y**, where *a* = 0 corresponds to the deafening case and *a* = 1 corresponds to the correct singing case.

### Experimental Methods

All experiments were performed in accordance with a protocol approved by the Duke University Institutional Animal Care and Use Committee.

### Surgery

Adult male birds (*n* = 8) were anesthetized with inhaled 1.5-2% isofluorane and placed on a head stereotax with a temperature-regulated bed. The incision was sterilized with alcohol, and a local anesthetic (bupivacaine 0.25%) was applied. Coordinates for the injections and lens implant were measured from the bifurcation of the midsagittal sinus. The coordinates used for targeting various auditory regions were as follows: CML, head angle of 43°, 1.8A, 1.95L, 0.7 and 0.9V; 2.0A, 2.05L 0.65 and 0.95V; 2.2A, 1.95L 0.7 and 0.9V. Birds were injected with 200nL of an AAV2/9 CAG GCaMP6s (Addgene, 100844-AAV9) viral construct on each site at a speed of 9nl/s using a Nanoject-II (Drummond Scientific). Immediately after the injections, we lowered a GRIN prism lens (Inscopix 1050-004601) held by a vacuum holder at a low speed (10um/s) centered around the injection sites and facing medially with the following coordinates: 1.5-2.5A, 2.1L, and 1.2V. After lowering, the lens was cemented to the skull and covered using body-double to prevent damage. Three weeks after the lens implantation we examined the field of view and cemented a baseplate to attach a 1-photon miniature microscope (nVista 3.0, Inscopix). If no ROIs were detected in the field of view during the baseplate implant or after recovery from anesthesia, the bird was excluded from the experiment. After two days of surgery recovery, the bird was attached to the microscope using a counterweight system to ensure normal behavior.

### Deafening

Adult male birds (*n* = 6) were deafened by bilaterally removing the cochlea. Birds were anesthetized with 20 mg/kg of ketamine and 10mg/kg xylazine and fixed on their side on a movable platform to facilitate access to the ear. The incision was sterilized with alcohol and anesthetized with bupivacaine 0.25%. The ear skin was cut using microscissors, and the tympanic membrane was punctured by a fine scalp. Then, the columella and the footplate were detached with forceps to reveal the oval window. A fine custom tungsten wire hook was introduced into the inner ear canal to remove the cochlea entirely. After surgery recovery (as measured by normal singing rates), the birds were re-attached to the miniscope to image the neurons after deafening. Before- and after-deafening comparisons were made between the day prior to deafening and the first day post-deafening when the bird resumed singing, typically between days 4-5 after deafening.

### Calcium imaging

Imaging data was acquired with a nVista 3.0 data acquisition board and the Inscopix imaging software at a frame rate of 12 Hz, using an LED power of 1-1.4mW/mm2. The z-plane was adjusted to maximize the number of focused ROIs and kept constant during the experiments. The acquisition board interfaced with custom Matlab code to monitor singing. The computer detected changes in amplitude when the bird started to sing to trigger the imaging system for sessions of 1 minute and record the audio data. The imaging files recorded within a day were curated by concatenating all the sessions that contained singing or playbacks together, spatially downsampling the videos, performing motion correction, and extracting the ROIs using CNMFe (74). The criteria for non-inclusion comprised sessions that had dropped frames or sessions that were accidentally triggered and did not contain song. The audio recordings containing song were manually labeled to determine song onsets and offsets. To avoid baseline contamination from cage noise or residual calcium associated with offset responses to prior song renditions, and considering the half-time decay kinetics of GCaMP6s (75), we only labeled songs that were preceded by at least 2 s of silence. Then, the extracted traces were aligned to the onset time stamps, z-scored, and averaged across trials. To generate the scatter plots in Fig. 4B, the activity was averaged over the first seconds of song.

### Cell registration

Cell registration pre- and post-deafening was carried out using CellReg (76). This algorithm used the spatial footprint matrices of the fields of view generated by the neuron extraction performed previously (74) to obtain a matrix of indices that corresponded to the same neurons across sessions. Specifically, CellReg translated and rotated the fields of view to match a reference session, yielding a new spatial footprint matrix in the same coordinate system. All neighboring cell pairs in close proximity were tested as follows: For every neighboring cell-pair, the algorithm calculated the distance between the center of mass (centroid distance) and the correlation coefficient between the spatial footprints. These metrics were calculated across sessions between nearest neighbors and far neighbors to generate a probability distribution of same-cell likelihood. These distributions were usually bimodal, with a threshold on distance or correlation to decleare cells the same optimized based on their separability. Centroid distance or spatial correlation models were selected on a case-by-case basis to maximize cell-registration yield. However, in both models, the threshold probability was set to p ¿ 0.9 to determine same-cell pairs and to minimize false-positive matches.

### Distorted Auditory Feedback

Custom software, EvTAF (53), was used to deliver white noise on 50% of randomly selected song renditions. Syllables were targeted based on matches against a template based on spectral frequencies, with matches triggering a 50-ms-long white noise playback emitted through a speaker at 80dB.

The neural responses were aligned to the onset of the first instance of white noise perturbation. The criteria for exclusion in this experiment included trials where the bird stopped singing immediately after the white noise perturbation; this prevented offset responses from being classified as error responses.

## Data, Materials, and Software Availability

The adult songs for training the sparse coding model and network models, as well as the correct songs in correct singing simulations, come from the control cases in a public dataset of segmented and labeled adult male zebra finch songs (47) (https://doi.org/10.18738/T8/SAWMUN). The juvenile calls and sound produced by the behaviors of juveniles for training the sparse coding model are randomly sampled from the recordings from a public dataset of juvenile zebra finch songs (48) (https://doi.org/10.7924/r4j38×43h). A cleaned and preprocessed version of these datasets that can be directly used for training the models in our study, as well as the code for simulating the models, running the analyses, and generating the figures in this work, can be found in https://github.com/gongziyida/vocal-error-network.git.

## ACKNOWLEDGMENTS

We are grateful for generous support from the Holland-Trice Foundation and the National Institutes of Health (grant nos. RF1DA056376, R01NS099288 and R01NS118424) in funding this research. We would also like to thank Dr. Kevin Franks for his insightful comments on this paper.

## Footnotes

† Note that Hebbian plasticity in the I→E connections is *effectively* anti-Hebbian, since the I→E connections are inhibitory.

* Certain memory modes can also have an effect on the shape of the error landscape. To isolate the effect of the components correlated with the tutor song patterns, we did not directly shuffle their entries.

