## Supplementary Information for "Correctness is its own reward: bootstrapping error signals in self-guided reinforcement learning"

Ziyi Gong[1], Fabiola Duarte[1] , Richard Mooney[1,2], John Pearson[1,3]

[1]Department of Neurobiology, Duke University, Durham, NC, USA
[2]Department of Cell Biology, Duke University, Durham, NC, USA
[3]Department of Electrical and Computer Engineering, Duke University, Durham, NC, USA

### Methods

#### Models

##### Simplified input for illustrating error landscape

The error landscape at a given moment in time is the transient neuronal activation around that moment. To be able to clearly visualize an error landscape in Fig.1A, we used a simplified auditory input pattern $\boldsymbol{\xi} \in \mathbb{R}^{N_E}$ with each element drawn from an i.i.d. Gaussian distribution, $\xi_i \sim \mathcal{N}(0, I_{3\times3})$, though the results do not depend on the mean and standard deviation as long as they are $\mathcal{O}(1)$. The premotor input is 1-dimensional. We would like to focus on a particular moment in time and thus set $r_H(t) = r^H_{\max}$ and $\mathbf{y}(t) = \boldsymbol{\xi}$. We then used the stationary population mean rates of the excitatory neurons at different auditory patterns to plot the error landscape.

##### Connectivity Matrices and EI Balance

The synaptic connections are represented by a set of weight matrices: $W_E$ denotes premotor→E connections and $J^{ab} \geq 0$ are connections from population $b$ to population $a$.

The values of the *initial* weights of existing synapses in the premotor→E connections are drawn from a log-normal distribution:

$$W_{ij}|_{c^W_{ij}=1} \sim \mathrm{LogNorm}(\mu_{W0}, \sigma^2_{W0}).$$

where $c^W_{ij} \in \{0, 1\}$ is a Bernoulli random variable with probability $P(c^W_{ij}) = \bar{c}_W$ and denotes whether the synapse from $r^H_j$ to $r^E_i$ exists. In the E→E and E→I→E models, $W_{ij}$ stays fixed, while in the feedforward and EI networks with premotor→E plasticity, it undergoes learning.

For the feedforward model, $J^{ab} = 0$, which reduces the equations to

$$\tau_E \frac{\mathrm{d}}{\mathrm{d}t}\mathbf{r}^E = -\mathbf{r}^E + \phi_E \left( W_E \mathbf{r}^H + \mathbf{y} \right). \tag{1}$$

For the EI network models, the *initial* weights obey

$$J^{ab}_{ij}|_{c^{Jab}_{ij}=1} \sim \mathrm{LogNorm}\left( \frac{J^{ab}_0}{\sqrt{N_b}}, \frac{(\gamma_{ab} J^{ab}_0)^2}{N_b} \right),$$

where $c_{ij}^{Jab} \in \{0, 1\}$ is again a Bernoulli random variable denoting whether the synapse from $r_j^b$ to $r_i^a$ exists, $J_0^{ab}$ controls the scales of the connections, and $\gamma_{ab}$ controls the variances. Other choices of $\gamma_{ab}$ with similar order of magnitude gave qualitatively similar results. For simplicity, we assume all recurrent weights have the same density, i.e. $P(c_{ij}^{JEE}) = P(c_{ij}^{JIE}) = P(c_{ij}^{JEI}) = P(c_{ij}^{JII}) = \bar{c}_J$.

To enforce EI balance, the $J_0^{ab}$ obey the EI balance condition

$$\frac{J_0^{EE}}{J_0^{IE}} < \frac{J_0^{EI}}{J_0^{II}}.$$

One can easily derived this inequality by noticing that the mean recurrent input to each neuron is $O(\sqrt{N\bar{c}})$ and must be set to zero:

$$\frac{I_{ext}}{\sqrt{N_E \bar{c}_J}} + J_0^{EE}\bar{r}_E - \sqrt{\frac{N_I}{N_E}} J_0^{EI}\bar{r}_I \approx O\left(\frac{1}{\sqrt{N_E\bar{c}_J}}\right) \approx 0$$
$$J_0^{IE}\bar{r}_E - \sqrt{\frac{N_I}{N_E}} J_0^{II}\bar{r}_I \approx O\left(\frac{1}{\sqrt{N_E\bar{c}_J}}\right) \approx 0,$$

where $I_{ext}$ is the total external input to each excitatory neuron and is assumed to be as strong as $O(\sqrt{N_E})$. Note that, in contrast with van Vreeswijk and Sompolinsky, 1998, the inhibitory population in our models does not receive strong external input. Unless otherwise mentioned, Table 1 summarizes the values of the parameters. Other values obeying the balance condition did not change the qualitative results.

For simplicity, $c_{ij}^W$ and $c_{ij}^{Jab}$ are fixed once the networks are initialized. That is, there is no birth or death of synapses during the simulations. Except for Fig. 3C (left), $P(c_{ij}^W) = 1$ for the feedforward and premotor→E models, and for E→E and E→I→E models, $P(c_{ij}^W) = 0.05$ for learning realistic input. In Fig. 3C (left), $P(c_{ij}^W)$ was varied from 0 to 1 to see its effect on the training in different models. Unless otherwise specified, $\bar{c}_J = 0.5$, though denser (up to 1) or sparser (0.3) recurrent connectivity did not show qualitatively different results.

#### Sparse Coding

Sparse coding theory postulates that neurons form an overcomplete set of basic sensory features, with only a small subset of neurons are activated for a particular sensory input Lewicki, 2002; Lewicki and Sejnowski, 2000; Olshausen and Field, 1996, 1997, 2004. Variants of sparse coding models have predicted key properties of sensory encoding Beyeler et al., 2019; Mukherjee et al., 2025; Olshausen and Field, 2004, including in primary visual cortex Olshausen and Field, 1996, 1997, primary and secondary auditory regions Lewicki, 2002; Mukherjee et al., 2025, and, pertinently to our work, in the caudo-medial nidopallium (NCM) of European starling Kozlov and Gentner, 2016, an auditory region containing neurons selectively tuned to birdsong.

We adapted a classic linear sparse coding model Olshausen and Field, 1996, 1997. Briefly, let $X \in \mathbb{R}^{K\times N}$ be a set of flattened spectrograms of bird song syllables, where $K$ is the number of samples and $N$ is the number of time bins multiplied by the number of frequency bins. We would like to find basis features $A \in \mathbb{R}^{L\times N}$, where $L$ is the number of features, and sparse coefficients (i.e. the neural responses) $S \in \mathbb{R}^{K\times L}$, such that

$$X = SA + \epsilon$$

where $\epsilon$ is some noise. In the original workOlshausen and Field, 1997, this was formulated as a probabilistic inference problem and approximated such that its solution for a training set $\tilde{X}$ can be found by repeating the following three steps until convergence of the basis $A^*$:

1. For a fixed $A$, find

$$S^* = \arg\min_S \frac{1}{N} \sum_{ij} (\tilde{X}_{ij} - (SA)_{ij})^2 + \frac{\lambda}{L} \sum_{ik} |S_{ik}|$$

2. For the optimized $S^*$, find

$$\hat{A}^* = \arg\min_A \sum_{ij} (\tilde{X}_{ij} - (S^*A)_{ij})^2$$

3. Normalize each basis, $A^*_{ij} = \hat{A}^*_{ij}/\sigma_{Aj}$, where $\sigma_{Aj}$ is the standard deviation of the $j$-th column vector.

We performed this optimization for a set $\tilde{X}$ comprising flattened spectrograms of normal syllables from adult male zebra finch songs Koch, 2024, immature vocalization Brudner, Samuel et al., 2022, and behavioral noise. The model targeted $L = 100$ basis elements, a highly overcomplete set for song, which is effectively low-dimensional at the syllable level ($d < 10$; Fig. S1A; Goffinet et al., 2021). Using the learned basis $A^*$, we then repeated Step 1 to find the sparse responses $S^*$ for normal syllables and syllables overlapped with 50-ms white noise, which were then used to train the vocal error model.

To map the $L$-dimensional sparse representations to $N_E > L$ excitatory neurons, we tried neighbor and random mappings, which resulted in qualitatively similar dynamics. For neighbor mapping, the $i$-th element of an $L$-dimensional sparse representation is the input to the excitatory neurons with indices from $(i-1) \times \lfloor N_E/L \rfloor + 1$ to $i \times \lfloor N_E/L \rfloor$. For random mapping, we generated a projection matrix $M \in \mathbb{R}^{N_E \times L}$ with elements each drawn i.i.d. from a unit normal distribution. While we found both approaches resulted in similar dynamics, the random mapping approach resulted in inputs that were more Gaussian and less sparse. Thus, to preserve input sparsity, we used neighbor mapping for this work.

#### Model Metrics

##### Percentage of Active Excitatory Neurons

To measure the sparseness of the population responses, we calculated the percentage of active excitatory neurons at every time step and then averaged over time window $[t_0, t_1]$, given by

$$P_{\text{active}} = \frac{1}{t_1 - t_0} \frac{1}{N_E} \sum_{t_0 < t < t_1} \sum_{i=1}^{N_E} H(r_i^E(t) - \theta_{\text{active}})$$

where $H(x) = 1$ if $x \geq 0$ and $H(x) = 0$ otherwise, and $\theta_{active} = 5$ Hz is the threshold chosen close to the spontaneous mean rate. Other values such as 3 Hz produced qualitatively similar results.

#### Cosine Similarity

We used cosine similarity to measure the relationship between the firing patterns across the neuron population and the auditory input patterns. For a population of $N$ neurons, the cosine similarity $S_c$ between the population vector at time $t$, $\mathbf{r}(t) = [r_1(t), r_2(t), ..., r_N(t)]^T$, and auditory pattern (either stationary or at some time step) $\mathbf{x} = [x_1, x_2, ..., x_N]^T$ is

$$S_c(\mathbf{r}(t), \mathbf{x}) = \frac{\mathbf{x}^T \mathbf{r}(t)}{\|\mathbf{x}\|_2 \|\mathbf{r}(t)\|_2},$$

where $\| \cdot \|_2$ denotes the Euclidean norm.

#### Quality of Cancellation

To measure the quality of cancellation, we compared how strongly the neurons represent the tutor song during correct singing trials before and after training. The strength of representation is measured by the cosine similarity described above. For each simulation, the quality of cancellation, averaged over the singing time window from song onset $t_0$ to song offset $t_1$, is then

$$Q_{\text{cancel}} = \frac{1}{t_1 - t_0} \sum_{t_0 < t < t_1} [S_c(\mathbf{r}_{\text{untrained}}(t), \mathbf{y}_{TS}) - S_c(\mathbf{r}_{\text{trained}}(t), \mathbf{y}_{TS})]$$

where the superscript $(i)$ denotes the $i$-th trial. The larger this quantity, the more effectively the cosine similarity between neural firing and the tutor song pattern is reduced. Negative values indicate that, on average, the representation of the tutor song is enhanced after training.

#### Normalized Neuronal Activity

Figs. 4, S2, S4, S6, and S7 contain normalized neuronal activity. For experimental data, the raw trial-averaged calcium activity of each ROI was first subtracted by the baseline (activity averaged over the 0.5-second window before singing onset). For Fig. 4B, the baseline-subtracted calcium activity was averaged over the 1-second time window after singing onset.

For Figs. 4C, S2D, S4B, S6D, and S7B, the processing is similar—trial averages subtracted by the baseline (0.1 s before song onset) and averaged over singing time window (1 s). A shorter window for the baseline is sufficient for our models and an computation-efficient choice, because our simulations have a much higher sampling rate (1000 Hz) than experiments (12Hz) and the transient dynamics is fast. Extending this window for baseline was found to have little effect. To get the trial averages, each trained model is simulated for 20 different initial conditions for Figs. 4, S4, S6, and S7, and for 50 different initial conditions for Fig. S2. Finally, for each model and each condition (correct versus perturbation or correct versus deafening), the averaged activities were divided by the standard deviation across neurons in the correct singing case (i.e., the x and y axes are rescaled by a common factor).

**Wasserstein Distance**

For Figs. 4D, S2E, S4C and S6E, Wasserstein distances were calculated to quantify the distances between the joint distributions of correct singing and perturbed (or deafened) singing responses from experiments and models. To remove the potential effect of different scales between models and experiments, for each condition and each source (experiment, feedforward, premotor→E, E→E, or E→I→E), both variables of the joint distribution were divided by the standard deviation across neurons in the correct singing case. The Wasserstein distances $l_w$ were then calculated between experiments and models, defined as

$$l_w(\mathbb{P}(\mathbf{x}), \mathbb{Q}(\mathbf{y})) := \inf_{\pi \in \Gamma(\mathbb{P}, \mathbb{Q})} \int \|\mathbf{x} - \mathbf{y}\|_2 \mathrm{d}\pi(\mathbf{x}, \mathbf{y}),$$

where $\mathbb{P}(\mathbf{x})$ and $\mathbb{Q}(\mathbf{y})$ are two probability distributions and $\Gamma(\mathbb{P}(\mathbf{x}), \mathbb{Q}(\mathbf{y}))$ denotes the set of distributions whose marginals are $\mathbb{P}(\mathbf{x})$ and $\mathbb{Q}(\mathbf{y})$. Intuitively, it measures how much effort is made minimally to convert one distribution to the other. We used SciPy's `wasserstein_distance_nd` function to carry out the calculation.

#### Bidirectional Sorting of Neuronal Activity

To visualize the sequential activation and sequential inhibition in Fig. S7, we developed a two-step process: First, the neurons were grouped into either excited or inhibited after song onset by comparing the mean activities before (500 ms for calcium imaging data and 100 ms for models) and after song onset (1500 ms for calcium imaging data and 1000 ms for models). Then, the excited group was sorted by the latency to the peak (maximum standardized activity) of each neuron, and the inhibited group was sorted by the latency to the dip (minimum standardized activity) of each neuron.

#### Connectivity Mode Analysis

To analyze the recurrent dynamics, we performed singular value decomposition (Fig. 5A) on the recurrent weight matrix over training time $t$,

$$\mathbf{J}_t = \begin{pmatrix} \mathbf{J}_t^{EE} & -|\mathbf{J}_t^{EI}| \\ \mathbf{J}_t^{IE} & -|\mathbf{J}^{II}| \end{pmatrix} = \mathbf{U}_t \mathrm{diag}(\mathbf{s}_t) \mathbf{V}_t^T, \tag{2}$$

where the columns of $\mathbf{U}$ and $\mathbf{V}$ are the left and right singular vectors, respectively, and s contains the corresponding singular values. Notice that we pull out the negative sign of the inhibitory weights explicitly to avoid confusion. The left singular vectors $\mathbf{U}_i$ ("modes") define the orthogonal directions to which the dynamics maps the population vector $\mathbf{h} = [h_1^E, ..., h_{N_E}^E, h_1^I, ..., h_{N_I}^I]^T$, where $h_i^a$ is the total input to the $i$-th neuron of population $a$. To see this, consider $\tau_E \approx \tau_I = \tau$ and rewrite Eqs. 4-5 as

$$\begin{aligned} \tau \frac{\mathrm{d}}{\mathrm{d}t} \mathbf{h} &= -\mathbf{h} + \mathbf{J}\mathbf{r} + \boldsymbol{\epsilon} = -\mathbf{h} + \mathbf{U}(\mathrm{diag}(\mathbf{s})\mathbf{V}^T\mathbf{r}) + \boldsymbol{\epsilon} \\ &\equiv -\mathbf{h} + \sum_i l_i \mathbf{U}_i + \boldsymbol{\epsilon}, \end{aligned} \tag{3}$$

where $\mathbf{h}$ is a concatenation of $\mathbf{h}^E$ and $\mathbf{h}^I$ and similarly for $\mathbf{r}$, and $l_i$ is the $i$-th element of $\mathrm{diag}(\mathbf{s})\mathbf{V}^T\mathbf{r}$.

#### Dissimilarity and Memory Encoding

The dissimilarity for each mode $i$ at the $n$-th rendition is defined as the maximum cosine similarity between itself and *any* mode at time 0:

$$\max_j |\mathbf{v}_i^T(n)\mathbf{v}_j(0)|,$$

where $\mathbf{v}$ is the corresponding left singular vector, and the 0-th rendition is the time prior to learning.

Similarly, to quantify the memory encoding for each mode $i$, we calculated

$$\max_k |\mathrm{Corr}(\mathbf{v}_i, \mathbf{y}_{TS,k})|,$$

the maximum correlation between that mode and $\mathbf{y}_{TS,k}$ the discrete pattern corresponding to the $k$-th syllable of the tutor song.

#### Convergence Time

We defined convergence time $\tau_c$ for our models as

$$\tau_c = \min\left\{t \left| \frac{x_t - x_{\min}}{x_{\max} - x_{\min}} > \theta_{\tau c}\right.\right\},$$

where $x_{\min}$ and $x_{\max}$ are the minimum and maximum of $\{x_t\}_t$, the parameter of the learning curve, and $\theta_{\tau c}$ is a threshold. We chose $\theta_{\tau c} = 0.8$. For learning in our data $t$ is measured in trials, such that $\delta t = 1$. In our analysis of network modes, we measured the convergence time for mean curves of the landscape and memory modes.

#### Perturbing the Modes

To probe the roles of different connectivity modes on the error codes represented by the excitatory neurons, we disrupted the excitatory patterns of the modes in several ways. For non-memory modes, i.e., those not correlated with the tutor song patterns, we directly perturbed their patterns using the following approaches:

1. shuffling all elements of the selected left singular vectors;
2. shuffling the first $N_E$ elements of the selected left singular vectors;
3. replacing the selected left singular vectors with patterns drawn from a zero-mean i.i.d. Gaussian distribution;
4. replacing the selected left singular vectors with zeros;
5. swapping the selected left singular vectors with the least significant left singular vectors.

All approaches produced qualitatively similar results. The first approach was used in Fig. 5, while the results from the others are shown in Fig. S10. Each perturbed left singular vector is scaled to have Euclidean norm equal to 1, except for Approach 4, which replaces the vectors with zeros.

For memory modes, we "de-memorized" them by subtracting the memory components from the left singular vectors. More specifically, denoting the excitatory part of the left singular vectors by $\mathbf{U}_E \in \mathbb{R}^{K \times N_E}$, where $K$ is the number of memory modes, the matrix of "de-memorized" vectors $\hat{\mathbf{U}}_E$ is

$$\hat{\mathbf{U}}_E = \mathbf{U}_E - \mathbf{U}_E \mathbf{M}^T \mathbf{M}$$

where the rows of $\mathbf{M} \in \mathbb{R}^{L \times N_E}$ are a set of orthonormal basis for $L$ syllable patterns ($\mathbf{M}\mathbf{M}^T = \mathbf{I}_{L \times L} \neq \mathbf{M}^T \mathbf{M}$). The "de-memorized" row vectors in $\hat{\mathbf{U}}_E$ are orthogonal to the basis vectors for the syllable patterns:

$$\begin{aligned}
\hat{\mathbf{U}}_E \mathbf{M}^T &= \mathbf{U}_E \mathbf{M}^T - \mathbf{U}_E \mathbf{M}^T \mathbf{M} \mathbf{M}^T \\
&= \mathbf{U}_E \mathbf{M}^T - \mathbf{U}_E \mathbf{M}^T \mathbf{I} \\
&= \mathbf{0}.
\end{aligned}$$

Finally, each row of $\hat{\mathbf{U}}_E$ is scaled to have Euclidean norm equal to 1.

### Supplementary Figures and Parameter Tables

| Symbol | Value | Definition |
|---|---|---|
| $N_E$ | 600 | Number of exc. neurons |
| $N_I$ | 150 | Number of inh. neurons |
| $N_H$ | 15 | Number of premotor neurons |
| $\tau_E$ | 30 | Exc. neuron time constant |
| $\tau_I$ | 10 | Inh. neuron time constant |
| $\sigma_\epsilon$ | 0.1 | Neuronal noise standard deviation |
| $r^E_{\text{max}}$ | 100 | Max. exc. neuron firing rate |
| $r^I_{\text{max}}$ | 100 | Max. inh. neuron firing rate |
| $\theta_E$ | 6 | Exc. neuron firing threshold (premotor→E / feedforward model) |
| $\theta_E$ | 0 | Exc. neuron firing threshold (E→I→E / E→E model) |
| $\theta_I$ | 0 | Inh. neuron firing threshold |
| $s_\phi$ | 2 | Neuronal activation function gain |
| $\bar{r}_{\text{burst}}$ | 150 | Mean peak firing rate of premotor burst |
| $\bar{\tau}_{\text{burst}}$ | 20 | Mean peak width of premotor burst |
| $\bar{\delta}_{\text{burst}}$ | 0 | Mean jittering of premotor burst |
| $\bar{c}_J$ | 0.5 | Recurrent connectivity probability |
| $J_0^{EE}$ | 0.1 | Scale parameter for E→E connections, $\mathbf{J}_{EE}$ |
| $J_0^{EI}$ | 0.17 | Scale parameter for I→E connections, $\mathbf{J}_{EI}$ |
| $J_0^{IE}$ | 0.1 | Scale parameter for E→I connections, $\mathbf{J}_{IE}$ |
| $J_0^{II}$ | 0.15 | Scale parameter for I→I connections, $\mathbf{J}_{II}$ |
| $\gamma$ | 0.1 | Recurrent connectivity variance parameter |
| $\mu_{W0}$ | 1/15 | Premotor projection mean |
| $\bar{c}_W$ | 1 | Premotor→E connectivity probability (premotor→E / feedforward model) |
| $\bar{c}_W$ | 0.05 | Premotor→E connectivity probability (E→I→E / E→E model) |
| $\tau_W$ | 10000 | Synaptic plasticity time constant |
| $\theta^E_{\text{active}}$ | 1.5 | Plasticity threshold for exc. neurons |
| $\theta^I_{\text{active}}$ | 5 | Plasticity threshold for inh. neurons |
| $\theta^H_{\text{active}}$ | 0 | Plasticity threshold for premotor neurons |
| $\Delta t_H$ | 10 | Plasticity time asymmetry (post - pre) for premotor→E synapses |
| $\Delta t_{EE}$ | 10 | Plasticity time asymmetry (post - pre) for E→E synapses |
| $\Delta t_{IE}$ | 10 | Plasticity time asymmetry (post - pre) for E→I synapses |
| $\Delta t_{EI}$ | 0 | Plasticity time asymmetry (post - pre) for I→E synapses |
| $\eta_H$ | -0.03 | Learning strength for premotor→E synapses (premotor→E model) |
| $\eta_{EE}$ | -0.05 | Learning strength for E→E synapses (E→E model) |
| $\eta_{EI}$ | 0.05 | Learning strength for I→E synapses (E→I→E model) |
| $\eta_{IE}$ | 0.006 | Learning strength for E→I synapses (E→I→E model) |

Table 1: **Model parameters in the main results, unless otherwise specified.**

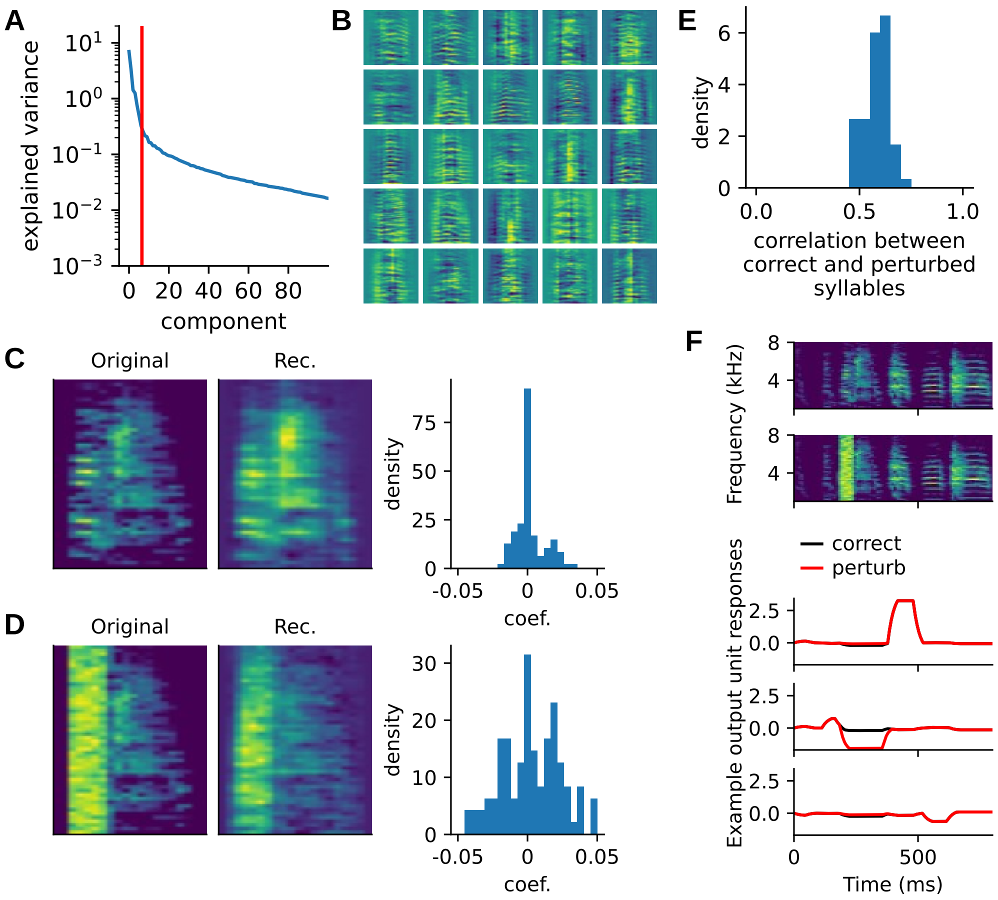


Figure S1: **Sparse coding of auditory input.** **(A)** The effective dimension (red) of tutor song syllable spectrograms, given by the participation ratio $d$ from the eigenvalues $\lambda_i$ of the covariance matrix between the flattened spectrograms. The participation ratio is given by $d = (\sum_i \lambda_i)^2/(\sum_i \lambda_i^2)$. Intuitively, if there are $K$ large eigenvalues $\lambda_1 \approx \lambda_2 \approx ... \approx \lambda_K \approx \bar{\lambda}$, $d \approx K^2\bar{\lambda}^2/(K\bar{\lambda}^2) = K$ (Gao et al., 2017). **(B)** Examples of learned basis elements. **(C-D)** Examples of reconstruction using the bases and corresponding coefficients (i.e., the responses of the sparse coding model; right column) for a normal syllable (C) and perturbed syllable (D). Right: histograms showing the distribution of coefficients for each reconstruction. **(E)** Responses from the sparse coding model for the perturbed syllables are still weakly correlated with the original syllables. **(F)** Syllable-specific neural responses over time. Top two rows show an example of correct song and perturbed song. Bottom three rows show three example auditory coding units.

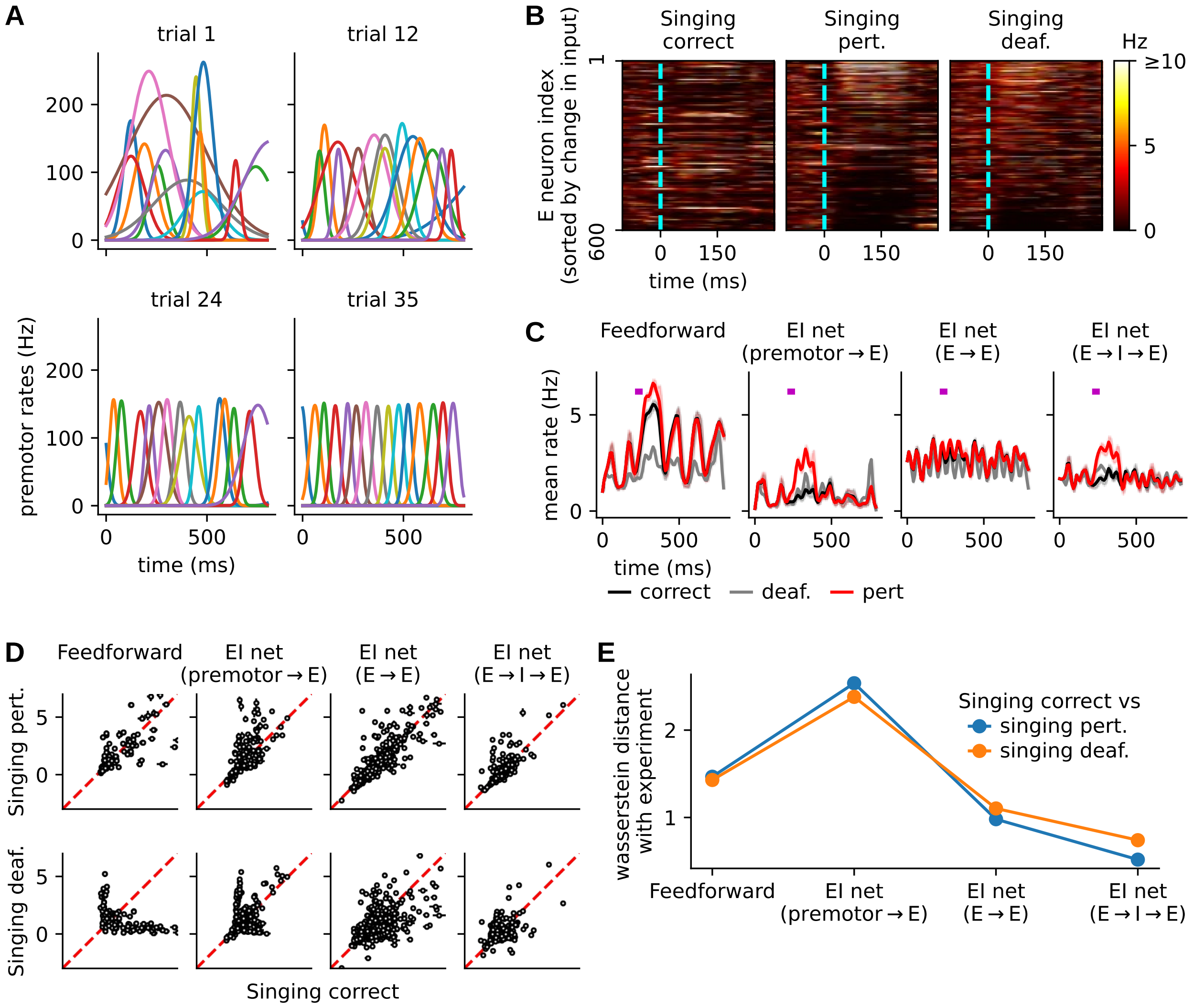


Figure S2: **Learning with gradually maturing premotor profiles.** With less temporally precise premotor input, the feedforward and premotor→E models are even more sensitive to firing thresholds. To see the full range of capabilities for each model, we chose excitatory thresholds $\theta_E = 6$ for the feedforward model, $\theta_E = 4$ for the premotor→E model, and $\theta_E = 0$ for the E→E and E→I→E models. **(A)** During training, the variances of the peak times and peak widths of premotor activities gradually reduce according to a flipped, smooth sigmoidal function of training time (rendition number; see Methods). **(B)** Trained models can represent the changes in auditory feedback compared to tutor song patterns as in Fig. 3D. The heatmaps show the responses of the E→I→E model, but the other models are qualitatively similar. **(C)** As in Fig. 3E, the population mean rates increase following perturbation in the premotor→E and E→I→E models, but not the E→E model. However, different from Fig. 3E, the mean rates in the feedforward model are not different between the correct and perturbation cases. **(D-E)** The distributions of trial-averaged responses in correct singing versus perturbed (top row) or deafened (bottom row) singing for different models trained with gradually regular premotor firing patterns **(D)**, and their Wasserstein distances with experiments **(E)**. The E→I→E model best reproduces experimental observations, in agreement with Fig. 4.

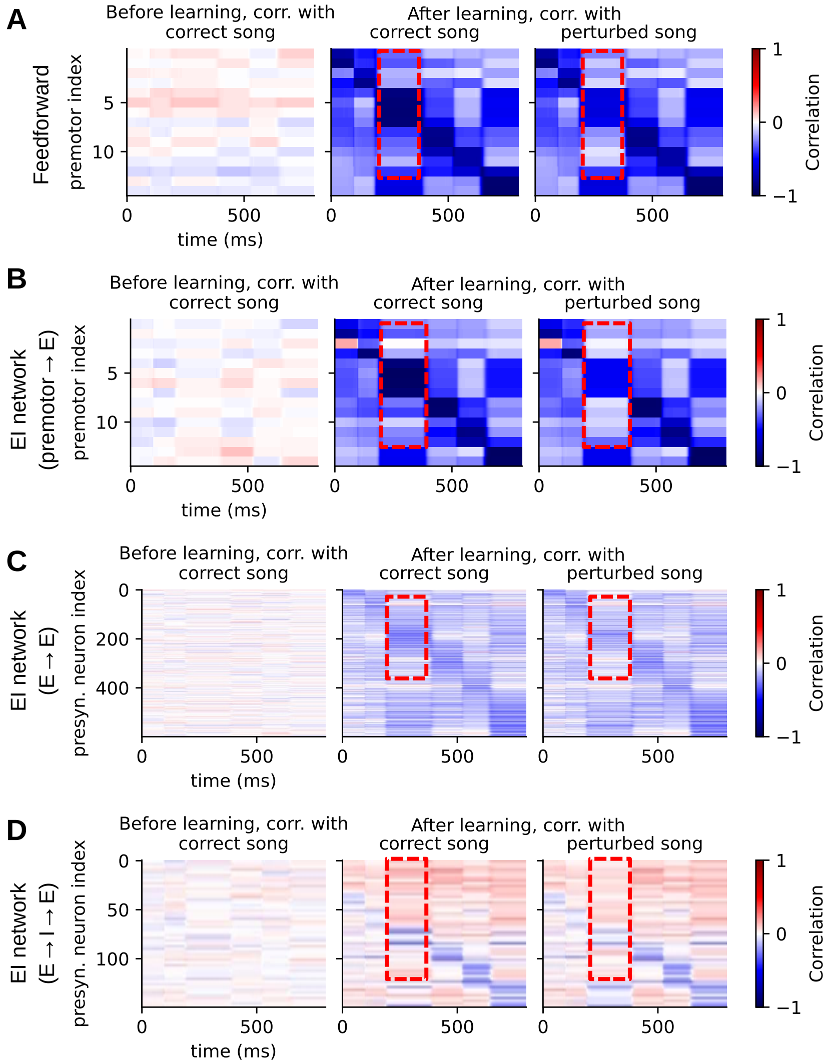


Figure S3: **Correlations between learned weights and sample-averaged tutor song auditory input patterns or perturbed patterns over time.** Left: correlations between the plastic weights and the sample-averaged target song $\mathbf{Y} \in \mathbb{R}^{T \times N_E}$ before training. The plastic weights for calculating the correlations are $\mathbf{W}_E \in \mathbb{R}^{N_E \times N_H}$ (premotor→E connections) in **(A-B)**, $\mathbf{J}_{EE} \in \mathbb{R}^{N_E \times N_E}$ (E→E connections) in **(C)**, and $\mathbf{J}_{EI} \in \mathbb{R}^{N_E \times N_I}$ (I→E connections) in **(D)**. The correlation is calculated between each column of the plastic weight matrix and each row of $\mathbf{Y}$. The middle and right panels are similar, but represent the correlations with the sample-averaged target song after training, and the sample-averaged perturbed song after training, respectively. Red boxes with dashed edges mark the time of perturbation and the synaptic weights that are less correlated with the perturbed patterns than the correct patterns.

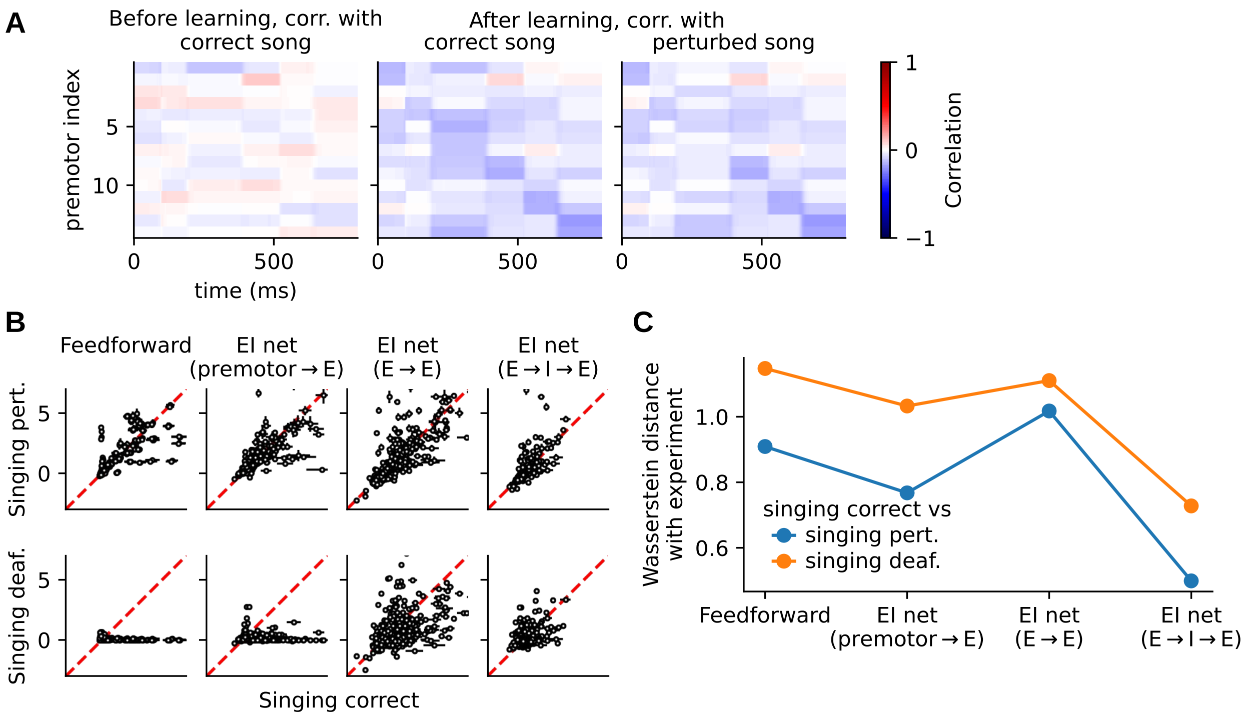


Figure S4: **Sparse premotor projections degrade error signaling in the feedforward and premotor→E models.** The density of premotor→E projection is 5% in this figure. Note that, in contrast, the premotor density is 100% (fully connected) in the feedforward and premotor→E models in the main results (cf. Fig. 3A-B, Fig. 3D-F, Fig. 4, Fig. 5, Fig. 6). E→E and E→I→E models were found to be insensitive to the premotor synapse density (Fig. 3C) and are re-simulated here just for comparisons. In the case of sparse premotor projections, **(A)** the learned weights and sample-averaged tutor song auditory input patterns or perturbed patterns over time are only weakly correlated (cf. Fig. S3). **(B-C)** Neurons are diversely modulated by perturbation and mostly silent in deafening, poorly matching the error responses observed in experiments (cf. Fig. 4).

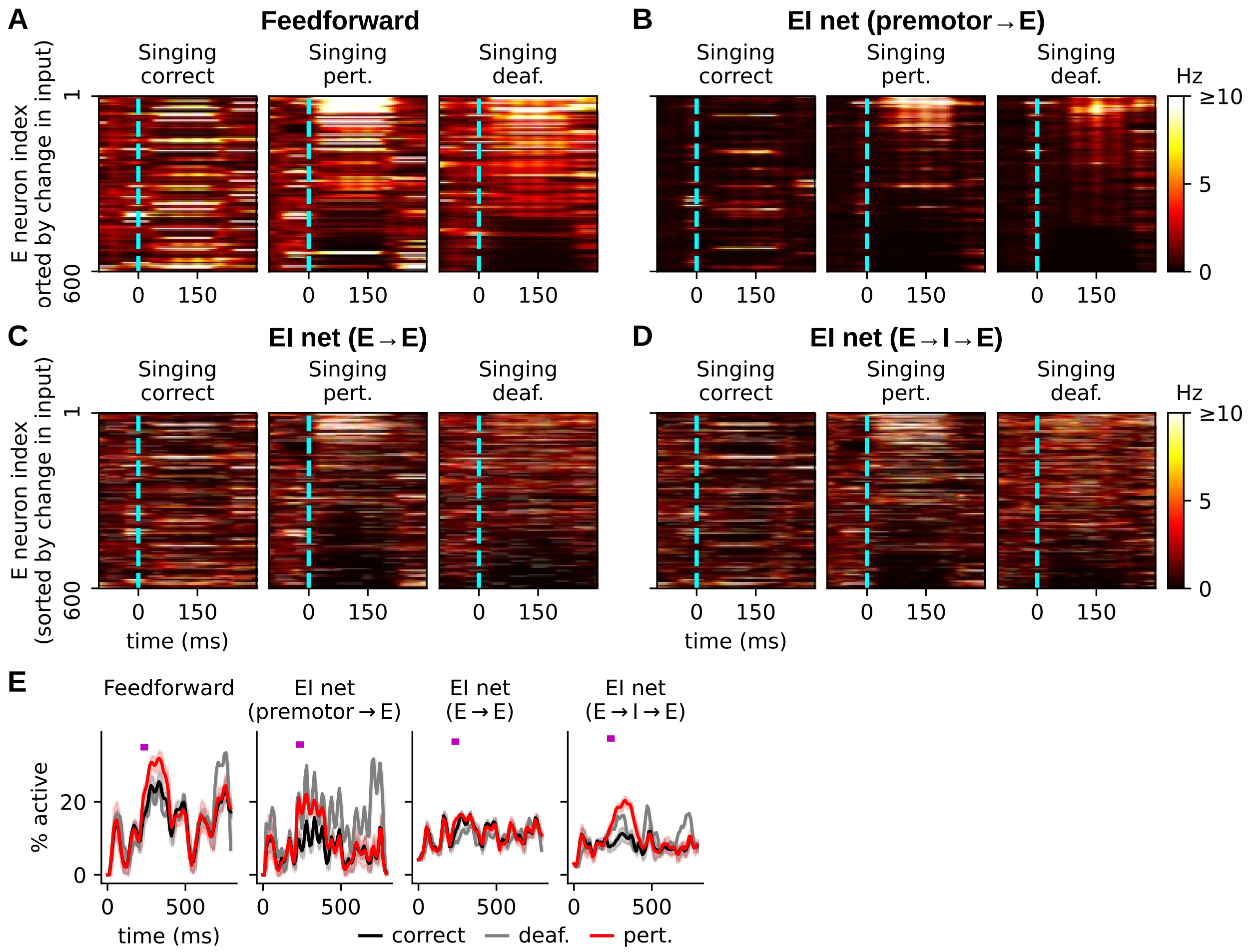


Figure S5: **Heterogeneous error responses in all four models.** **(A-D)** Learned singing responses under normal practice and perturbation in all four models, plotted in the same way as Fig. 3D. Note that Fig. 3D is for the E→I→E model and another example trial is included in **(D)** in this figure for comparisons between models. While the responses are qualitatively similar across models, EI networks display more temporally variable and sparser responses than the feedforward model. **(E)** Percentage of active neurons over time for different models. The purple bar in each subplot indicates the 50-ms white noise perturbation to the auditory feedback of the bird's own song. Following the perturbation, the feedforward, premotor→E, and E→I→ models increase activation density for around 100 ms. The activation density for the E→E model, however, remains stable during white noise perturbation.

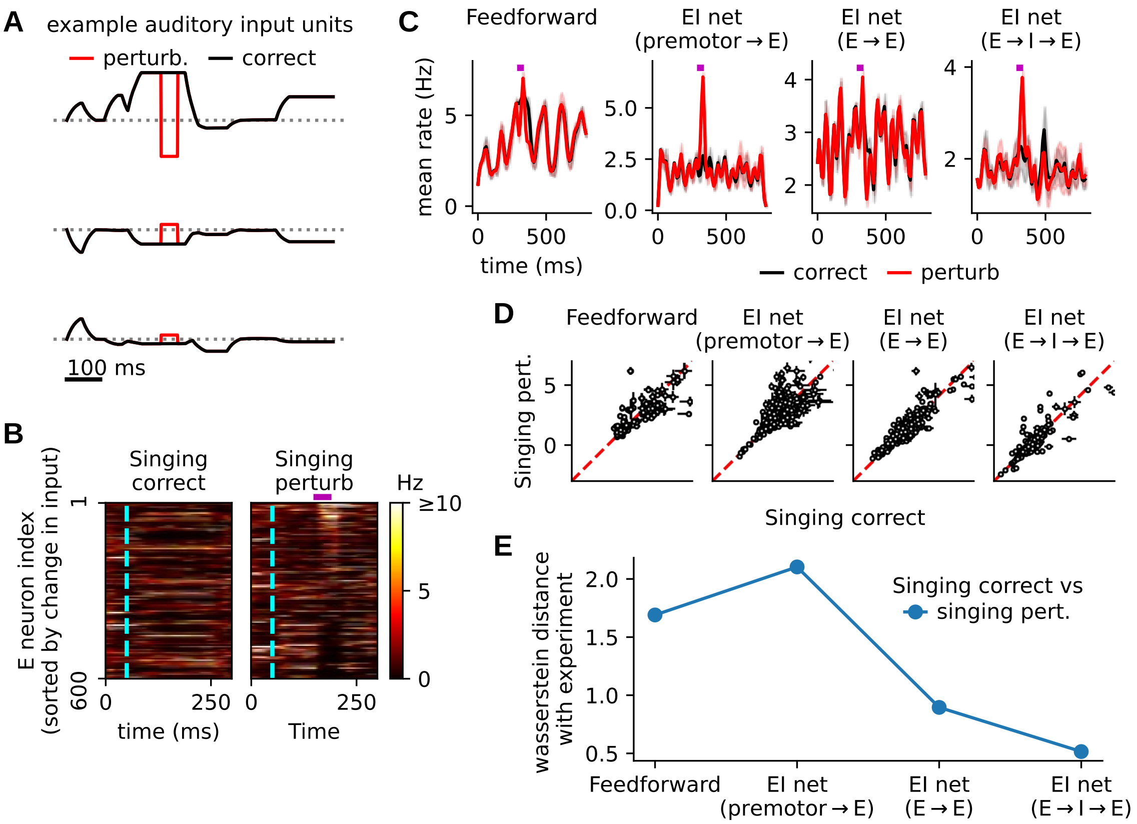


Figure S6: **Models are able to signal perturbation at sub-syllabic level.** In the main text, models were tested with white noise perturbation superimposed over the audios of the targeted syllables, which were then encoded by a sparse coding model and extended over time, thus changing the whole neural encoding of the targeted syllables even if the perturbations were much shorter (50 ms) than the syllables ($\sim$150 ms). Here, this experiment tests the error signaling ability when the neural encodings are perturbed for only 50-ms duration and remain unchanged outside the perturbation window. **(A)** Three example input units for the correct (black) and perturbed (red) cases. Correct auditory inputs are the sparse representations of the tutor song recordings, as for the experiments in Figs. 3 and 4. The perturbed inputs are constructed by substituting a 50-ms chunk of the correct inputs with a random white noise pattern whose elements are drawn from $\mathcal{N}(0, \sigma^2_{correct})$ where $\sigma^2_{correct}$ is the variance of the correct inputs (red steps in the curves). The 50-ms sub-syllabic perturbation occurs in the middle of a $\sim$150-ms long syllable. **(B)** Learned error responses in the E$\to$I$\to$E model. Cyan dashed lines indicate the start of the perturbed syllable, and the purple horizontal bar indicates the duration of the 50-ms sub-syllabic perturbation. Only shortly (50-100 ms) after the 50-ms sub-syllabic perturbation, the neurons corresponding to larger change in the auditory input patterns have higher firing rates. The other models are qualitatively similar to the E$\to$I$\to$E model. **(C)** Similar to the perturbation at the syllable level (Fig. 3E), the population mean rates in the models except for the E$\to$E model are increased by the sub-syllabic perturbation, but for a much shorter duration. **(D-E)** In agreement with experiments (Fig. 4), sub-syllabic perturbation activates a sparse set of neurons, and the E$\to$I$\to$E model best reproduces experimental observations (cf. Fig. 4.)

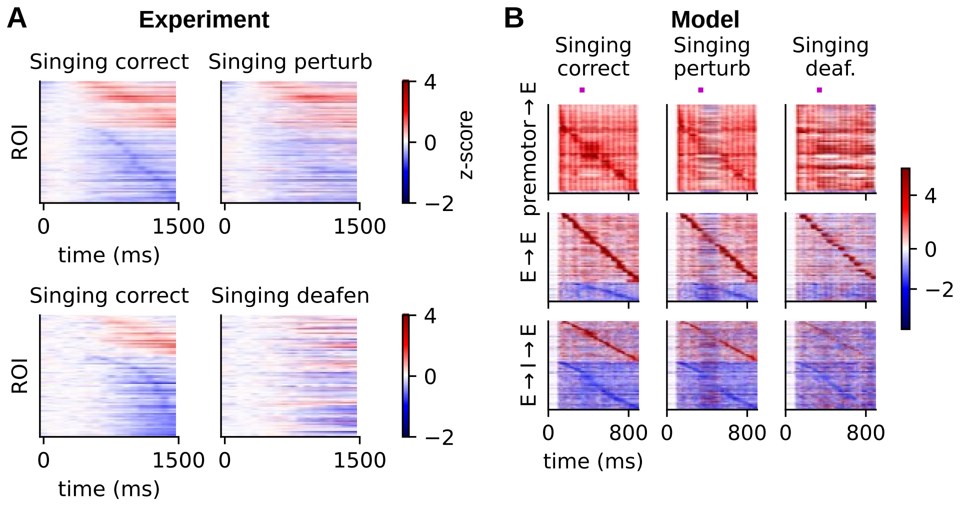


Figure S7: **The E→E and E→I→E models can reproduce both the sequential activation and suppression observed in experiments.** **(A)** Top, normalized calcium activity in CM during the correct (left) and white noise perturbation (right) cases. Bottom, normalized calcium activity in CM during the correct (left) and deafening (right) cases. In the correct case, neurons display sequential activation or suppression. The order of the sequential activity in the correct singing case is disrupted by perturbation or deafening. **(B)** Normalized neuron firing rates for the premotor→E (top), E→E (middle) and E→I→E (bottom) models, during the correct (left), perturbed (middle), and deafened (right) singing cases. The feedforward model is qualitatively similar to the premotor→E model. All models are able to reproduce the sequential activation observed in experiments during correct singing. However, only the E→E and E→I→E models can also produce the sequential suppression. The disruption of sequential activity during perturbation is more obvious in the premotor→E and E→I→E models than in the E→E model. Each row of **(A-B)** was standardized using the mean and variance calculated over the displayed time window. In both panels, the neurons in the perturbed singing cases were sorted in the same orders as those in the corresponding correct singing cases.

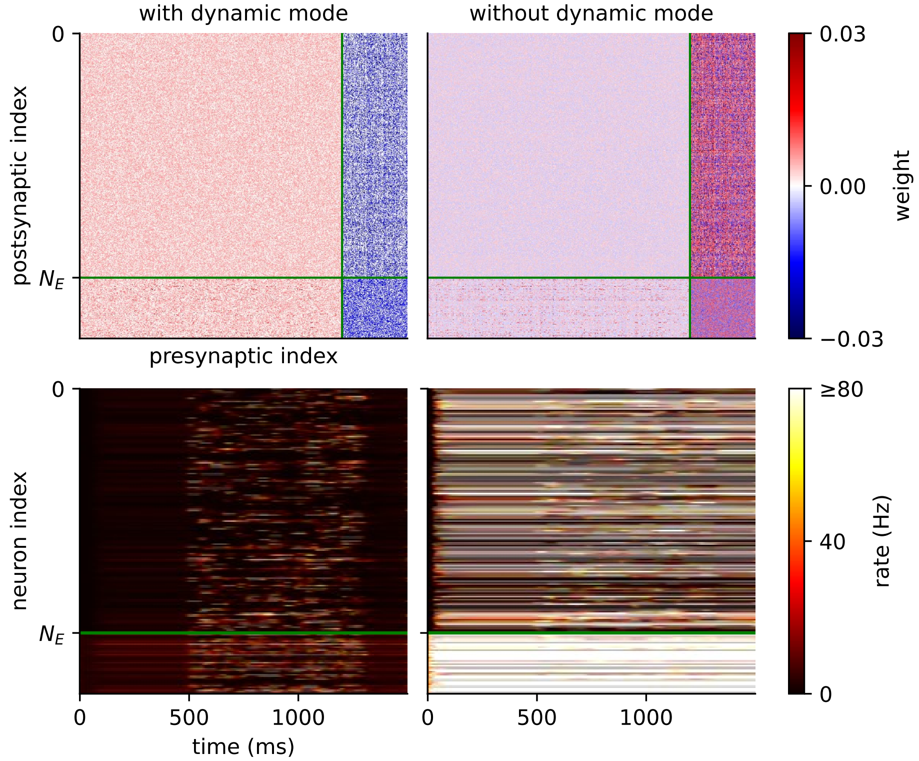


Figure S8: **Removing the dynamic mode in the E→I→E model drastically changes the connectivity and breaks E-I balance.** Top left: original connectivity matrix **J** post-learning. Excitatory and inhibitory synaptic weights are shown in red and blue colors, respectively. Green lines mark the boundary between excitatory neurons and inhibitory neurons. Top right: connectivity matrix post-learning, but with the dynamic mode removed. The removal was done by setting the mode to zero, but the results were qualitatively the same when other perturbation methods were used (see Methods). Bottom left: sparse and mild neuronal firing with the original connectivity weights. Bottom right: dense and hyper-active neuronal firing when the dynamic mode is removed.

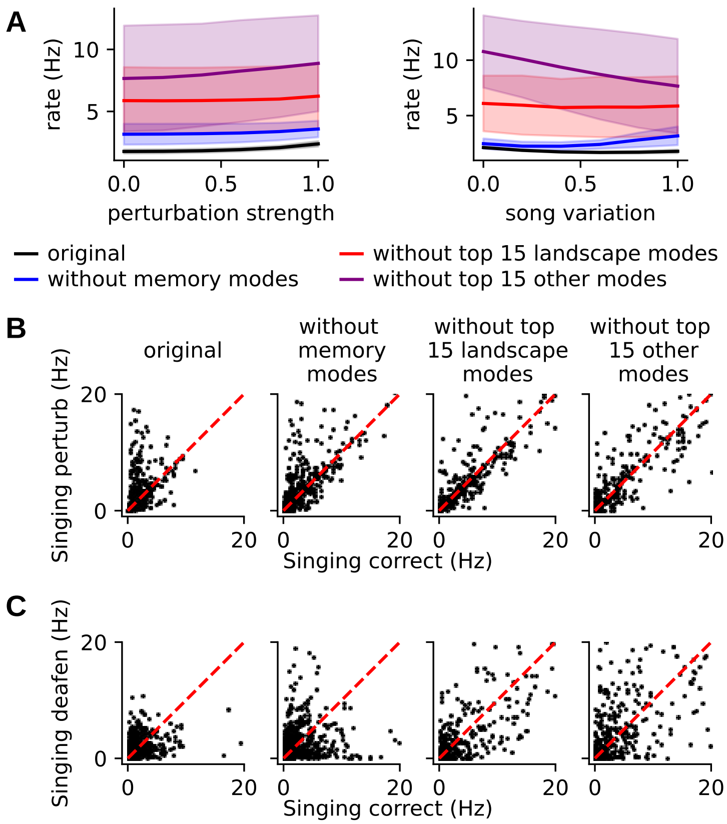


Figure S9: **Connectivity mode perturbations significantly change population activity.** **(A)** Excitatory population mean rates as functions of perturbation strength (left) and song scaling (right) for the original trained models (black); models with top 15 landscape modes removed (red), with memory modes removed (blue); and with top 15 non-memory, non- landscape modes removed (purple). Compared to the original models, all three interventions significantly changed the mean firing rates for every choice of perturbation strength and song scaling ($p < 10^{-4}$, two-sided Wilcoxon rank-sum test). **(B-C)** Distributions of trial-averaged excitatory rates between the correct singing case and perturbed **(B)** or deafened **(C)** conditions for the original trained models (leftmost column) and the three interventions (three right columns).

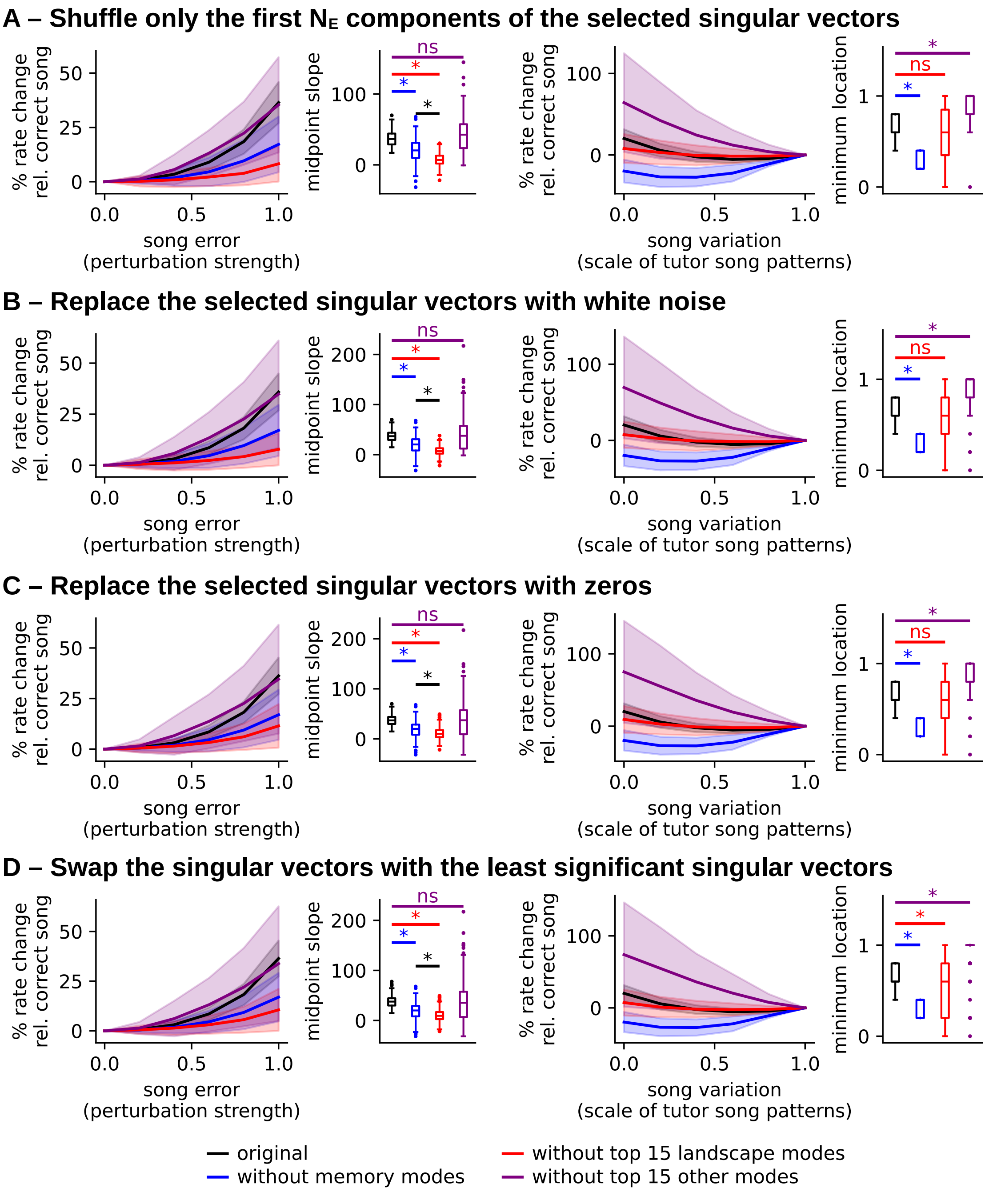


Figure S10: **Perturbing the connectivity modes using alternative approaches.** Each row shows the results from one alternative approach (see Methods) to perturbing the connectivity modes, plotted in the same way as in Fig. 5F-G. All results qualitatively agree with Fig. 5 that perturbing the landscape modes most strongly flattens the error landscape, and only perturbing the memory modes significantly moves the landscape minimum towards the silent input.

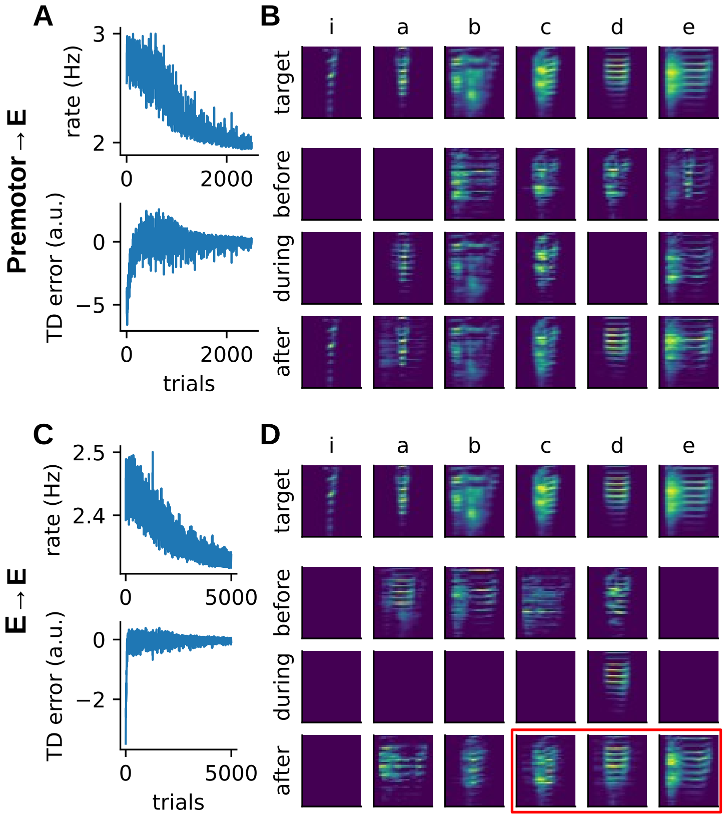


Figure S11: **Error codes produced by the premotor→E and E→E models can also be used to train a motor policy**. Top row, premotor→E model; bottom row, E→E model. **(A)** and **(C)** as Fig. 6B, and **(B)** and **(D)** as Fig. 6C. The red box in **(D)** indicates the generated syllables that match the target syllables.

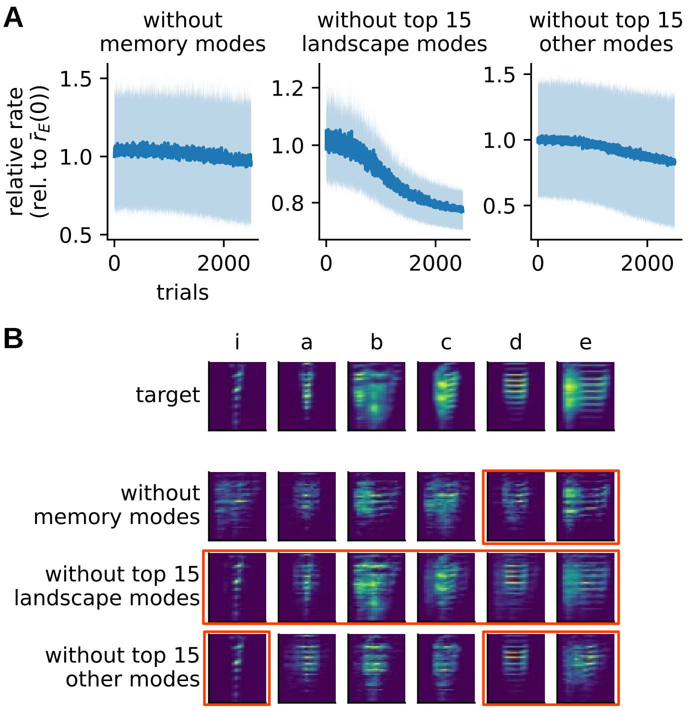


Figure S12: **Syllable generation via RL is impaired when using the error codes from the E→I→E models with perturbed connectivity modes.** **(A)** The excitatory population rate (error signal) barely decreased over training trials when the memory modes were removed, but decreased robustly in the absence of the top 15 landscape modes. Removal of the top 15 non-memory, non-landscape modes resulted in a slightly decreasing error signal on average, though with very high trial-to-trial variance. Solid curves and the vertical widths of the shaded areas represent the mean and std, respectively, over $n = 12$ random initializations. **(B)** Tutor syllable templates (top row), and mean generated syllables after RL using the E→I→E models with different perturbation conditions of the connectivity modes (bottom three rows). The mean was taken over $n = 12$ random initializations of training. Taking the median does not qualitative change the results. Red boxes indicate the relatively well-matched syllables. Compared with Fig. 6, the production of many syllables is negatively affected, and more syllables are incorrect in the case with perturbed memory modes than in the other two scenarios.
